# Inferring Toll-Like Receptor induced epitope subunit vaccine candidate against SARS-CoV-2: A Reverse Vaccinology approach

**DOI:** 10.1101/2020.12.24.424322

**Authors:** Ananya Nayak, Ayushman Gadnayak, Maheswata Sahoo, Shivarudrappa B Bhairappanavar, Bidyutprava Rout, Jatindra Nath Mohanty, Jayashankar Das

## Abstract

Toll-Like Receptors (TLRs) are a group of Pattern Recognition Receptors (PRRs) which bind to the exogenous pathogen associated molecular patterns (PAMPs) like other PRRs; hence the main function is to sense the harmness and mediate the innate immune response to pathogens. TLRs play an important role in innate immune responses to infection. The host has evolved to use other TLR and PAMP agonists as agents to stimulate a protective inflammatory immune response against infection. Because only a small number of doses are given, TLR agonists appear to have greater potential and fewer safety concerns than other uses as vaccine adjuvants. In the present days, development of peptides targeting immune response can be approved for survival in biological monitoring systems before vaccine exposures. Peptide vaccines are easy to synthesize, more stable and relatively safe. In addition, production of peptides becomes simple, easily reproducible, fast and cost effective. Getting vaccinated against Covid-19, which has become a pandemic in the human population, is the most practical way to control the outbreak. The new coronavirus does not contain a drug or vaccine to prevent it from spreading to humans. To getting a proper vaccine candidate against the novel coronavirus, the present study used the reverse vaccinology approach by using a complete set of SARS-CoV-2 proteins; such as: Spike, Envelope, Nucleocapsid, Membrane, NSPs, and ORFs to extract the antigenic elements that produce B-cell, T-cell and IFN positive epitopes. These epitopes with precise binding to the Toll-Like receptors (1-10) have developed epitope based vaccine candidates. We have prioritized a set of epitopes based on their antigenicity, allergenicity, sequence conservation and projected population coverage world-wide. The selected epitopes were employed for in-silico docking interactions with Toll-Like receptors and molecular dynamic simulation confirmed the stability of the vaccine candidates resulting epitope of spike proteins with both the TLR 7 and 8 shows the best binding affinity. We believe that this ideal epitope vaccine candidate could enhance the immune response of the host and reduce the reinfection risk.

## Introduction

COVID-19 infection and its high transmission rates have spurred research and industry to develop effective vaccines and drugs. However, since then there has been no effective vaccine or drug to fight this infection. There is still no suitable strategy for therapeutic treatment, drugs, or vaccines [1]. Current clinical data indicate that the severity is due to cytokine storms and immunopathogenesis leading to inflammation in the lungs, especially in the alveolar tissue. Given the importance of the disease COVID-19 to biology, the TLR-SARS-CoV-2 interaction appears to be an appropriate target for the concept of a suitable therapeutic strategy against the pandemic [2, 3, 4].

Nowadays, the incidence of serious diseases is increasing day by day all over the world due to the lack of adequate therapy and vaccine development [5, 6, 7, 8]. Vaccination against Covid-19 which has become a pandemic in the human population is the most practical way to control the outbreak [9, 10, 11]. The novel coronavirus does not contain a drug or vaccine to prevent it from spreading to humans. In this regard, we are concerned about the need to develop a vaccine candidate against this deadly virus. Immunizations are an essential part of primary health care. Vaccines are the prevention and control of infectious disease outbreaks. They contribute to global health security and play an important role against antibiotic resistance. However, traditional vaccine production methods take decades to overcome pathogens and antigens, disease and immunity. To support this, reverse vaccinology is an approach that reduces the time and cost of identifying vaccine candidates whose traditional methods have failed [12, 13, 14].

Two decades ago, classical or traditional methods of identifying protein targets were used to develop recombinant vaccines against microbial diseases [15, 16]. Traditional methods for developing vaccines involving whole organisms or large proteins result in unnecessary exposure to antigens and an increased likelihood of allergic reactions. This problem can be overcome by using peptide vaccines containing short fragments of immunogenic peptides that can cause a strong and targeted immune response, thus avoiding the risk of allergic reactions. Recent advances in computational biology have opened new doors for the design of effective silicon vaccines. This study used an immuno-informatics approach to obtain a peptide vaccine candidate against SARS-CoV-2 [17].

Peptide based vaccines are essentially an in-silico approach that does not require in-vitro culture [18]. Peptide preparation is simple, easy to prepare, faster and less expensive for solid phase peptide synthesis. In considering the novelty of a peptide vaccine, it is most important to choose the right peptide [19, 20]. When selecting the right peptide, the antigen plays an important role in the effectiveness of the vaccine for example: the pathogen life-cycle, molecular-pathogenesis including host entry, and pathogen survival. Therefore, vaccines can be developed to trigger an immune response that can trigger pathogen invasion [21, 22, 23]. The main mechanism of peptide vaccines is based on chemical methods of synthesizing B-cells and T-cell epitopes, which can trigger specific immune responses. B-cell epitopes bind to T-cell epitopes to make them immunogenic [24, 25]. In this study, we used the Immune Epitope and Resource Analysis Database (IEDB) to catalog the data available for other coronaviruses [26, 27, 28]. Parallel bioinformatics prediction a priori identifies potential B and T cell epitopes for SARS-CoV-2. This prognosis can facilitate the effective design of high priority vaccines against this virus [29, 30, 31].

Various types of immune cells, such as dendritic cells, macrophages, monocytes, myelin-expressing T lymphocytes, B cells, initiated microglia and responsive astrocytes, are important for the pathogenesis of infection [32, 33, 34]. Toll Like Receptors (TLRs) are a group of pattern recognition receptors (PRRs) which binds exogenous pathogen associated molecular patterns (PAMPs) hence involved in pathogen identification and protection of hosts located on the surface or endosomes of various cell types such as dendritic cells, macrophages, B cells, T cells, epithelial cells, oligodendrocytes, astrocytes and endothelial cells [35], which plays an important role in triggering the innate immune response by producing IFN Type I inflammatory cytokines and various mediators [36, 37]. TLR agonists and antagonists have been suggested as compounds with broad therapeutic potential against various respiratory diseases associated with antiviral drugs and vaccine adjuvants [37, 38]. Among all types, TLR3 receiving double-stranded RNA (dsRNA), TLR7 and 8 viral transmits single-stranded RNA and is therefore likely involved in SARS-CoV-2 clearing. The main protein for myeloid 88 differentiation response (MyD88) is an adapter that binds TLRs to downstream molecules [38]. Activation of TLR via MyD88 and TRIF-dependent pathways induces nuclear translocation of transcription factors NF-κB −3 and IRF-7 with the development of innate anti-inflammatory cytokines (IL-1, IL-6, TNF-α) and type I. IFN-α / β, which is important for antiviral responses [39].

Currently, immune responsive target peptides can be developed and approved for survival in biological monitoring systems before vaccines are available for exposure and practical studies [40]. We believe this ideal epitope vaccine can increase immune response and further reduce the risk of reinfection by increasing the immunogenicity of the host [41, 42]. In this study, it was hypothesized that advanced computer screening could identify peptides suitable for subunit vaccine preparation. We used the complete set of SARS-CoV-2 proteins to extract antigenic elements that produce B-cell and T-cell immunity. By integrating these peptides, we have developed epitope-based vaccine candidates with precise binding to the Toll like Receptors 7 and 8 (TLR-7 and 8) for an effective immune response and with the lowest possible toxicity and hypersensitivity. Since this method has been validated experimentally with constructs against other pathogenic species, we assume that the proposed structural formulation has the potential to produce an effective immune response against the coronavirus. Wet laboratory researchers are expected to validate our design with the hope of providing protection for healthy communities from the COVID-19 pandemic(Figure 1).

**FIGURE 1:**
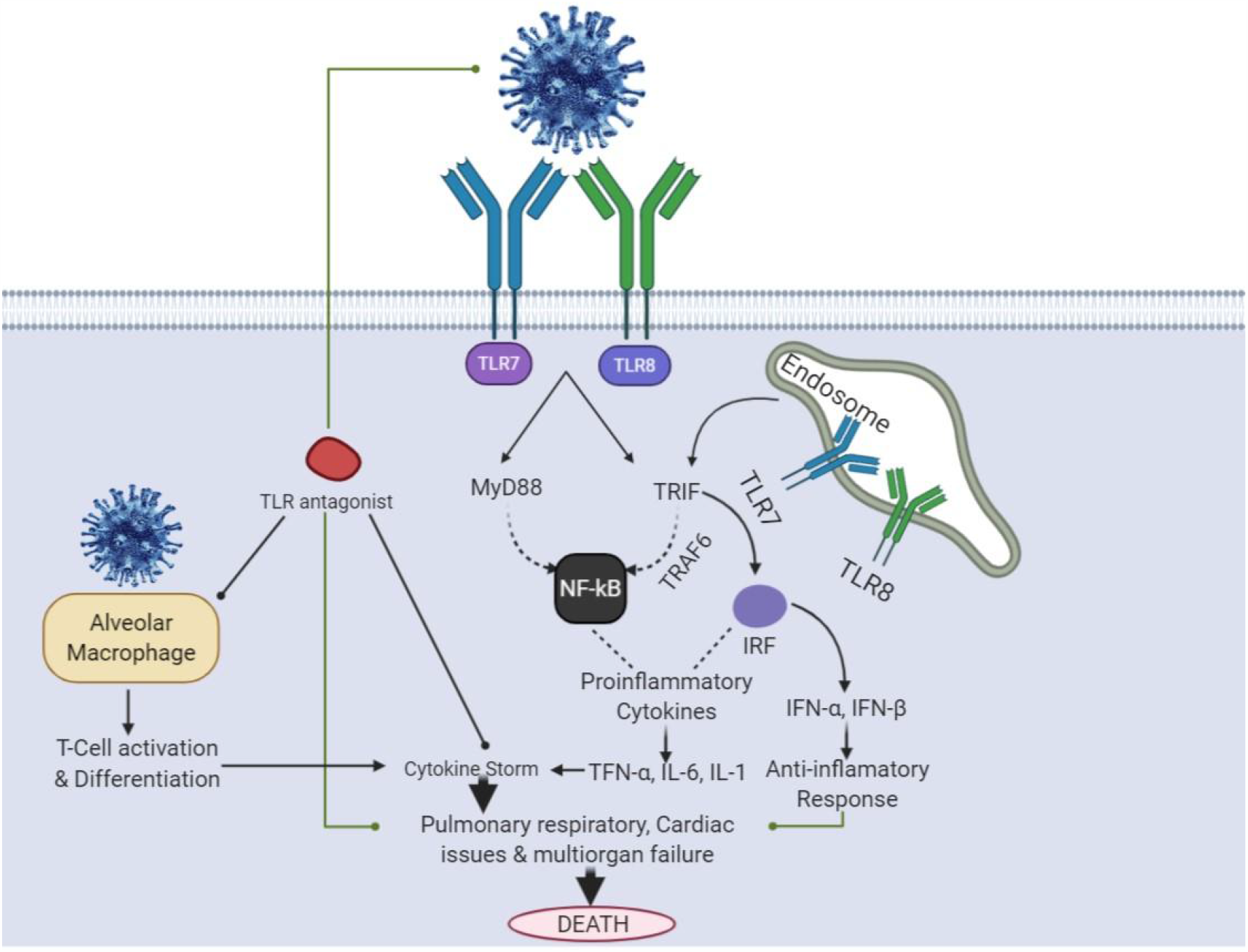
Toll-Like Receptors (TLRs) as a potential target for therapeutic intervention against SARS-CoV-2. NFκB, nuclear factor κB; IL, interleukin; IFN, interferon.

## MATERIALS AND METHODS

### Retrieval of Protein Sequences and selection

Whole proteomes of the SARS-CoV-2, encoding all proteins were retrieved from NCBI [43] and reviewed proteins were retrieved from the Universal Protein (UniProt) [44] database in FASTA format.Pathogen mediated core proteins were determined by using **CD-HIT suite** [45] that withdraw proteins sharing sequence identity of 60%.

### Antigenicity prediction

By setting the default parameters, all protein sequences were sent to **VaxiJenv2**.**0** to find out the highest antigenic protein. **VaxiJenv2**.**0** [46] is designed to predict the most potent subunit vaccine candidate with an accuracy of 70-90%. All antigenic proteins with the highest antigenicity scores were selected for further evaluation.

### Subcellular localization

Recognition of immunogenic proteins by immune cells to stimulate the immune response is one of the crucial factors for the selection of good vaccine candidates, which can be studied from the subcellular localization. Therefore, **CELLO2GO** [47] was deployed to estimate the protein sequences with the highest antigenic score.

### Linear B-cell epitope prediction

Based on their spatial structure, the B-cell epitopes can be ordered in two types; a linear or continuous and another was conformational or discontinuous. The linear B-cell epitopes was predicted using the Immune Epitope Database (IEDB) [48]. Surface accessibility, hydrophilicity and flexibility properties are also important. Therefore different methods were implemented respectively along with the default parameters of IEDB to predict these properties, We have predicted the linear B-cell epitope using Bepipred linear epitope prediction method. The location of linear B-cell epitopes was predicted by Bepipred using a combination of a Hidden Markov Model and a propensity scale method. The residues with scores above the threshold (default value is 0.35) were considered to be a part of an epitope.

### Helper T-cell (HTL) Epitope Prediction

The requisite of T cell receptors to epitopes complexed with major histocompatibility complex class II (MHC) molecule can lead to T cell activation; To predict MHC class II restricted HTL epitopes, protein sequences were subjected to NetMHCII Pan 3.1 server [49] with a threshold value of 0.5% and subject to a 2% value for strong binding peptides (SB) and weakly binding peptides (WB) respectively, to determine the binding affinity of the MHC-II epitope and allele. Here a firmly binding epitope with the maximum number of binding HLA-DR, DP and DQ was selected as a possible candidate for the epitope.

### Prediction of Cytotoxic T-lymphocyte(CTL)

Prediction for CTL epitopes is crucial for consistent vaccine design. Therefore, the amino acid sequence constituting the CTL epitopes of selected protein was predicted using **NetCTL**.**1**.**2** [50] server along with the default parameters.

### IFN Gamma (γ**) Epitope Prediction**

Gamma interferon (IFN-γ) is a cytokine that is essential for innate and adaptive immunity and, apart from stimulating natural killer cells and neutrophils, acts as the primary activator of macrophages. For innate and humoral immunity, IFN-γ plays an important role in antitumor, antiviral, and immunoregulatory activities. Therefore, IFN-γ-inducing epitopes were important for the design of a potential multi-epitope vaccine [19]. For the designed peptide, 15-mer IFN-γ epitopes were predicted using the **IFNepitope server** [51]. For the adjuvant and the main vaccine peptide IFN-γ epitope predictions were carried out separately, due to the restrictions in the no. of residues that can be used for prediction by the server.

### Conservancy analysis

In order to check the conservancy of the epitopes, they were evaluated among all the strains present globally by using the epitope conservancy analysis tool at the **IEDB** [48] analysis resource.

### Allergenicity assessment

The allergenicity of all the conserved epitopes were taken to be analysed using **AllergenF pv**.**1**.**0 server** [52], which is based on the amino acids in the protein sequences in data sets were described by five E-descriptors.

### 3D Structure of epitopes

The 3D structures of all the non-allergenic and 100% conserved epitopes were modelled using **I-TASSER** [53], **ORION**: web server [54] and **PEP-FOLD 3** server [55].

### Homology Modelling and structure analysis of Target proteins

For homology modelling, all the TLR proteins of human (1-10) were retrieved from UniProt and modelled using the **Phyre2** server [56], followed by refinement of the modelled proteins, by using **3DRefine** server [57]. Model validation is a significant advance to identify possible errors in rough structures and to analyze the quality of models during the refinement process. The refined model quality assessment of the model has been done by generating **Ramachandran Plot** by using **Schrodinger server andPROSA** [58]. The ProSA server assessed the general quality of the protein models as a z-score. At the point when the predicted model’s z-scores are outside the typical range of primary proteins, it demonstrates an inaccurate structure. The 3D structure of protein was visualized using PyMol [59].

Based upon iterative threading assembly and simulation methods, **I-TASSER server33** [60] generated five 3D models for the individual protein sequence and ranked all the models based on their C-scores. C-score values measure similarity between the query and template based on the significance of threading template alignment and the query coverage parameters. The Ramachandran plot analysis of refined structure revealed the percentage of residues were located in most favoured region, most allowed and generously allowed region, while also showed the residues were in disallowed region. However, ProSA-Web calculated Z-score of −4.52 indicating the model was not in the range of native protein conformation. Furthermore, the ModFold6 server was used to evaluate the overall quality of the model.

### Human Receptor Proteins for Molecular Docking Studies

For molecular docking study, to ensure the interaction between human receptor proteins include 10 TLR of human and our predicted non allergenic, 100% conserved CTL epitopes using the **Autodock 4**.**2**.**6** [61]. The protein ligand complex showing best binding affinity will be further proceed for MD-Simulation in order to check the stability.

### Molecular dynamics simulations

Molecular dynamics simulations were carried out to check the stability of the anchored epitope receptor protein complexes using the **GROMACS v2016**.**3** software [62]. For each complex, a production simulation of 50 ns at 350 K and a pressure of 1 bar was obtained after protocols for minimal energy savings and for balancing the solvation system had been carried out. In addition, path analysis was performed to check for H-transition and Root Mean Square Deviation (RMSD).

### Population Coverage

Population coverages for sets of T cell epitopes were computed using the tool provided by the Immune Epitope Database (IEDB) [48]. This tool uses the distribution of MHC alleles (with at least 4-digit resolution, e.g., A*02:01) within a defined population to estimate the population coverage for a set of T cell epitopes.

## Results

### Protein targets

A total 192720 protein sequence of SARS-CoV-2, encoding all types of proteins was retrieved from NCBI and 14 reviewed proteins were retrieved from the UniProt (Universal Protein) database in FASTA format. Then, CD-HIT analysis was performed to identify the pathogen core proteins by “a clustering approach that relies on multiple alignments” and got 31 clusters having proteins sharing identity of 60% (Table s1).

### Antigenicity prediction

In-order to detect the most potent antigenic protein, all the protein sequences were analyzed by the VaxiJen server, which were also retrieved and revealed the most potent antigenic protein. Among 31 proteins, 26 proteins were identified as the highest potential antigenic protein that is 0.6123 of total prediction score as well (Table 1).

**TABLE 1:**
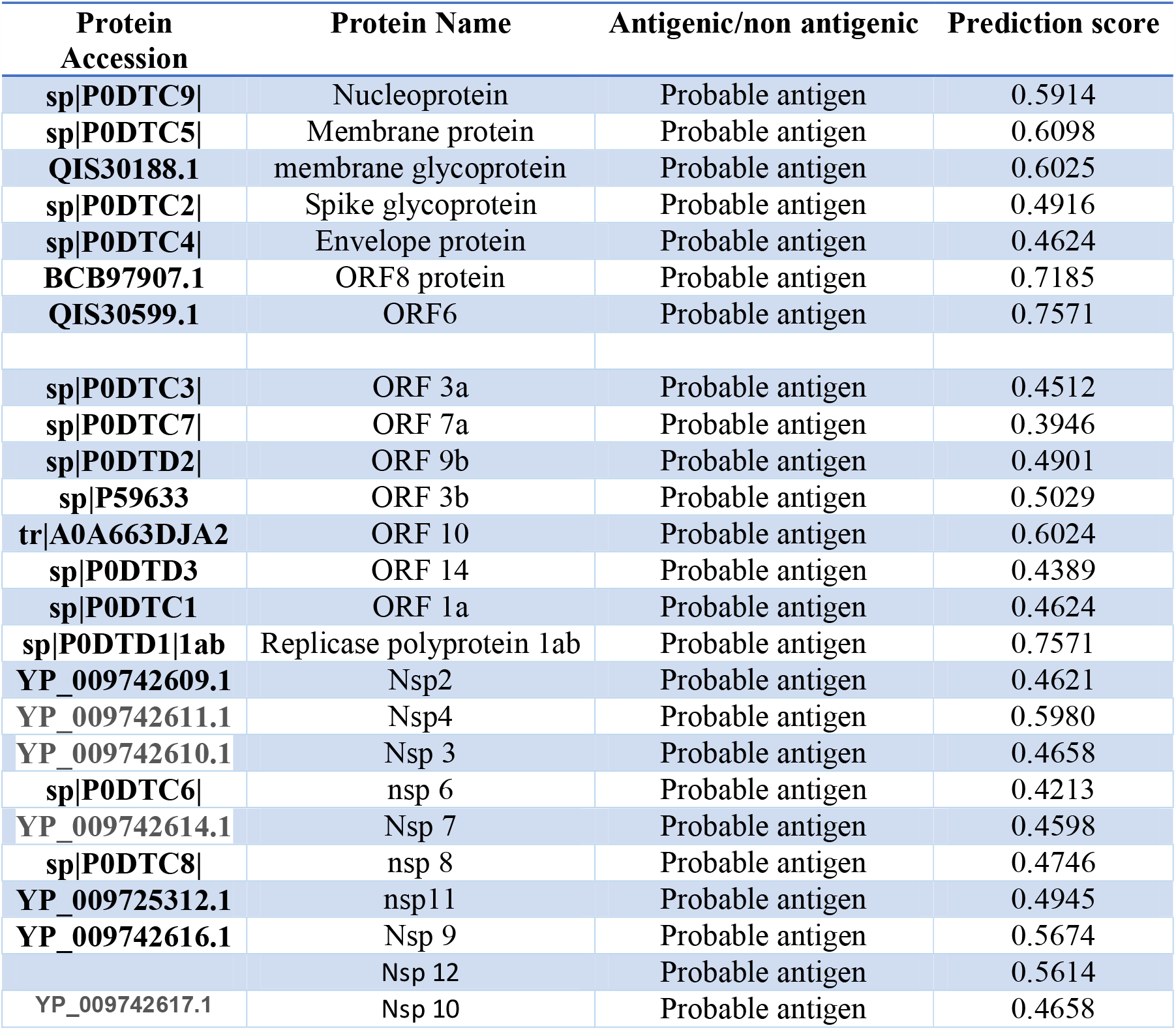
Antigenicity prediction of all structural, non-structural and accessory proteins from SARS-CoV-2 with protein ID and prediction score.

### Sub-cellular localization

As we all believed that the sub-cellular location of the protein plays an important role in determining its function. In this study we identified sub-cellular location of all our 26 most antigenic proteins from which 8 proteins were found in extracellular part of a eukaryote model likewise 18 proteins were found in plasma membrane, two were found in cytoplasm, one in mitochondria and three were found in chloroplast. All show diverse molecular functionality like RNA binding, structural molecule activity, transmembrane transporter activity, protein binding (Figure 2).

**FIGURE 2:**
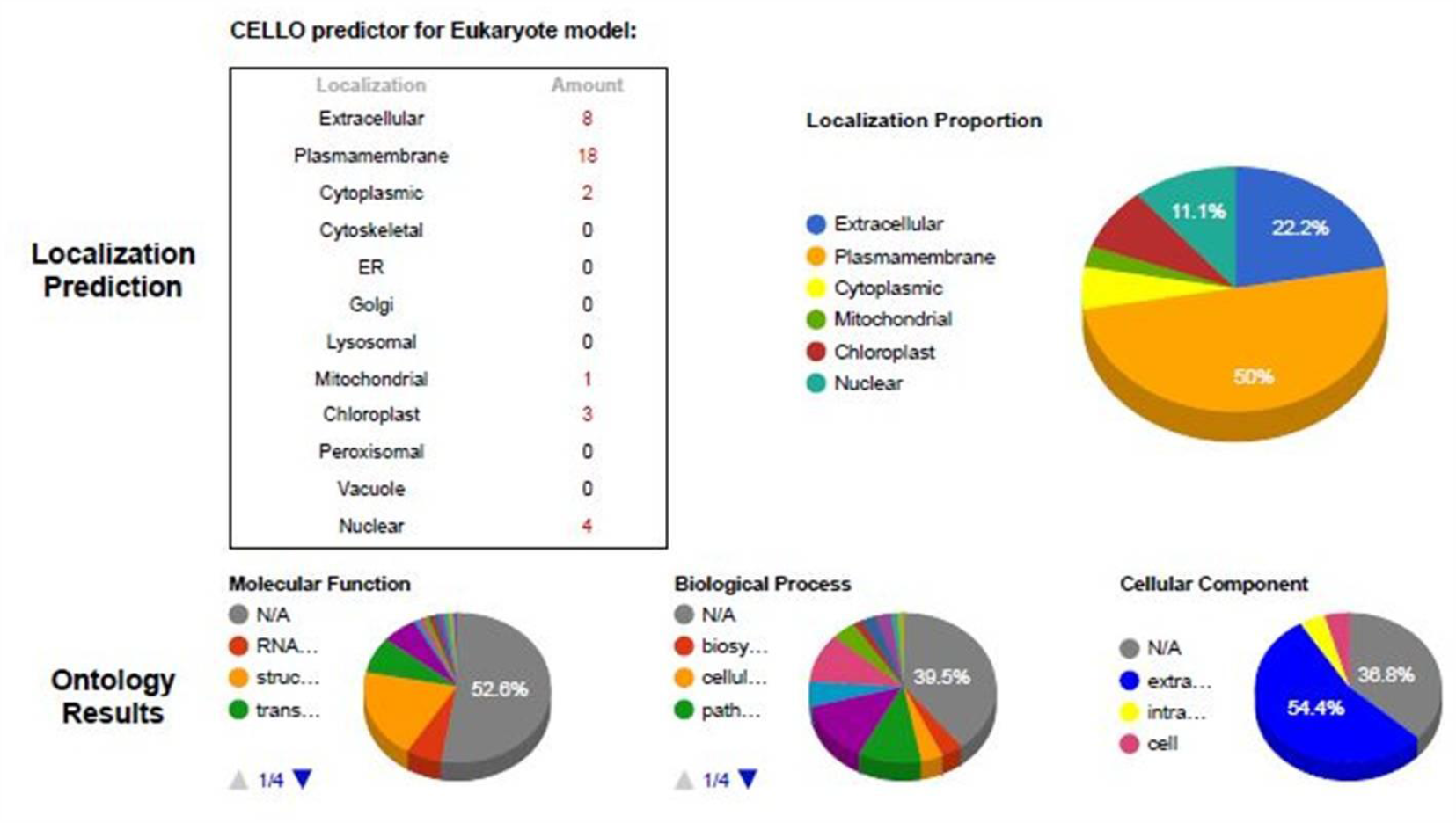
The number of proteins shown in each subcellular location represents the number of ‘locative proteins’. The dataset comprises 38 proteins distributed in 14 subcellular locations.

### Primary and secondary antigenic protein structure determination

In this study, we analyzed the most significant antigenic protein to comprehend the secondary structure as well as their physiochemical characteristics property. We obtained protein lengths as well as their molecular weights in theoretical Isoelectric point (PI) and in Daltons in result. The instability index (II) was computed, which showed that the protein sequences are all stable. Also we calculated the positively charged and negatively charged residues. The atoms are found having H, C, N, O, and S with the analysis of amino-acid composition of these proteins. The striking average of hydropathicity (GRAVY) was obtained to be negative by the calculation of aliphatic index, which indicated the protein hydrophilic nature. So the protein is liable to have enhanced interaction with other proteins. The study of protein secondary structure found the dominancy of random coils with extended strand of α-helix and β turns (Table 2).

**TABLE 2:**
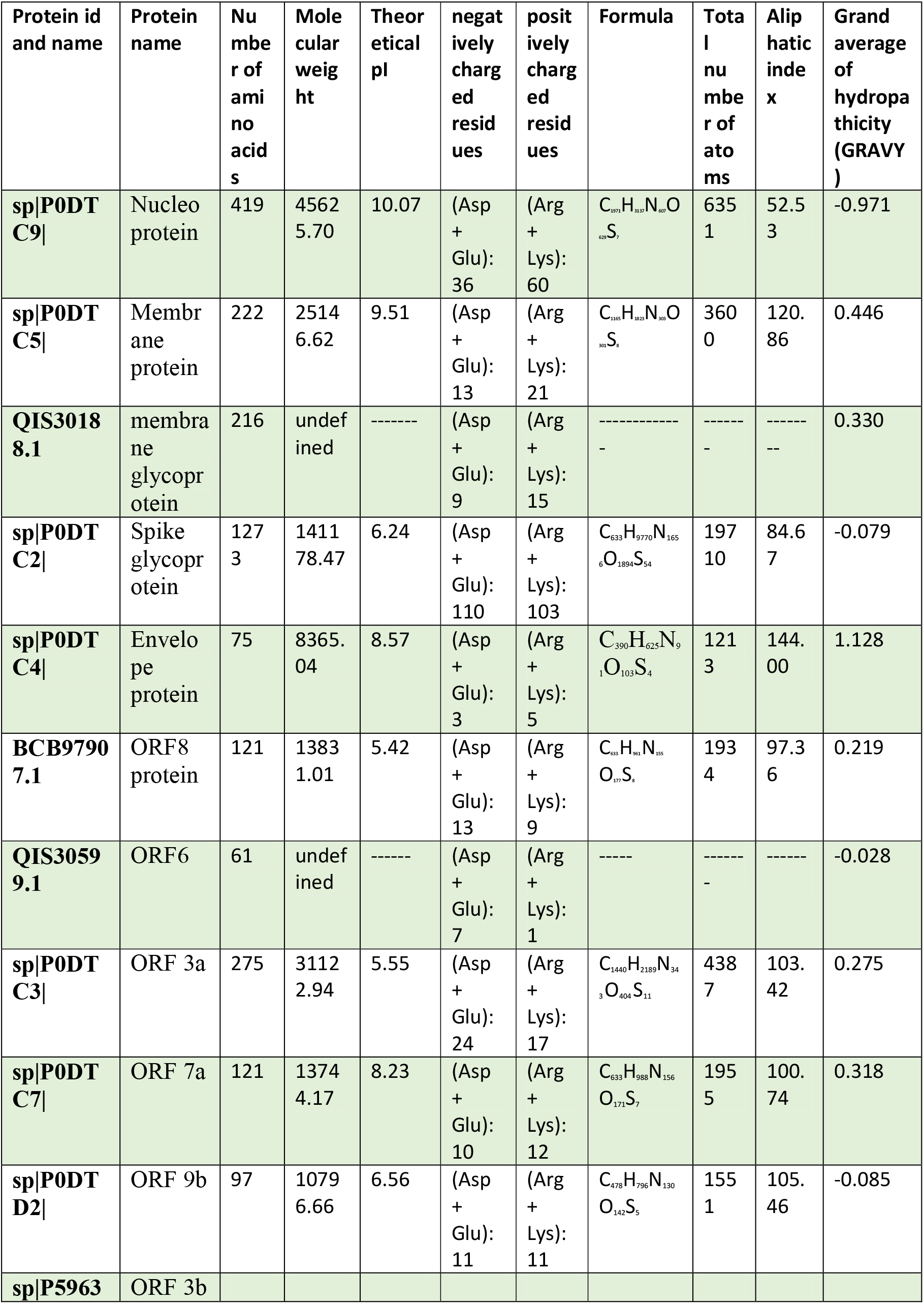
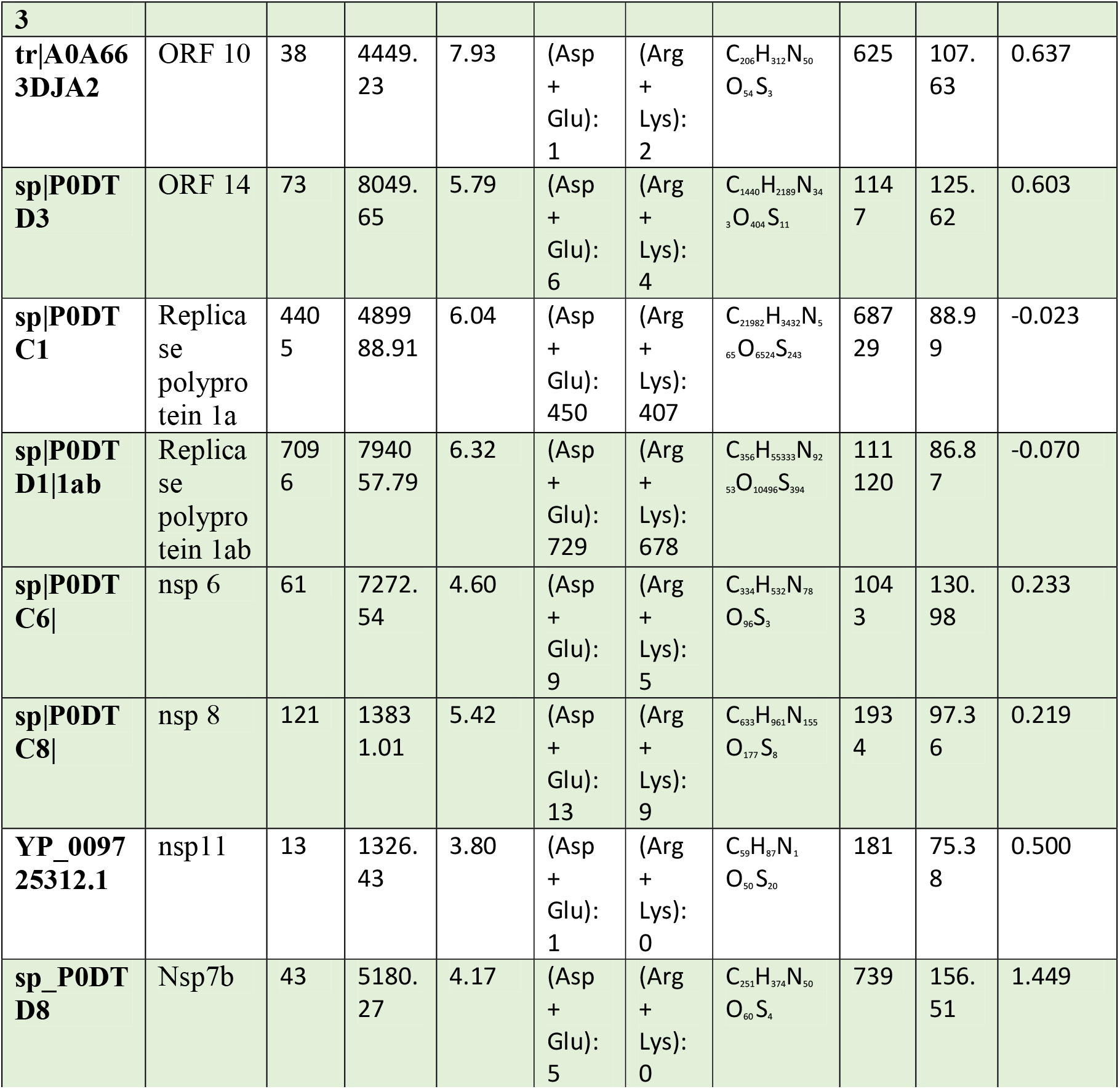
Different physio-chemical properties of SARS-CoV-2 proteins with Formula and GRAVY score.

### Linear B-cell epitopes detection

The most significant steps in epitope based vaccine design are the detection & characterization of B-cell epitopes in target antigen. The antigenic peptides prediction was done by IEDB (Immune Epitope Database) analysis resource derived from the Kolaskar and Tongaonkar’s method, where determination of antigenic epitopes were by analyzing the physiochemical properties of aminoacid. The output 26 antigenic proteins are found having 430 antigenic peptides in the amino acid range of 5-20 length. We also predicted the maximum residual score for every amino acid in this analysis. Out of that some residues were predicted as conformational with CTL epitopes that can be taken as valuable candidates for peptide based vaccine development. Besides, the result revealed about the average antigenic propensity score of the predicted epitope was 1.002 and the maximum and minimum score was 1.237 and 0.550. In the X and Y-axis the graphical representation of sequence position and antigenic propensity were shown which represents the predicted antigenic residues.

### T-Cell Epitope Prediction

T cells when bound to the MHC molecule, it recognizes epitopes. So epitopes prediction only is possible after computing their MHC-binding profile. The epitopes binding prediction to MHC I is more precise than to MHC II as the molecular interactions between epitopes & MHC I and MHC II complexes are different. In our IEDB analysis study, we identified 765 predicted CD8+ T cell epitopes which were exclusive and established coming from different *T. cruzi* known protein regions found in the *T. cruzi* H > 0.5 masked proteome.At first, 8068 HLA class I epitopes were anticipated inside glycoproteins of SARS-COV-2. Investigation based on percentile rank sifted 1150 peptide epitopes by setting the boundary of binding affinity>0.3. Every one of them had an extensive binding affinity for the 12 superfamily alleles. These epitopes are chosen alongside their highlights and respective binding alleles. Activation of T cells is possible by binding of T cell receptors with the epitope complex along with MHC class II molecules. The binding affinities of MHC II and respective epitopes are determined by the server NetMHCIIpan having threshold values 2% and 0.5% for weak binding peptides and strong binding peptides respectively. In our case we selected putative epitopes candidates’ characteristics by the strong binding epitopes with more number of binding affinity with HLA-DP, DR and DQ allele. From all HLA (DP, DQ and DR) 30,711 epitopes were predicted. We have taken all (765) the epitopes below 50% affinity for further study. In Table 3 we have provided selected epitopes with their respective alleles.

**TABLE 3:**
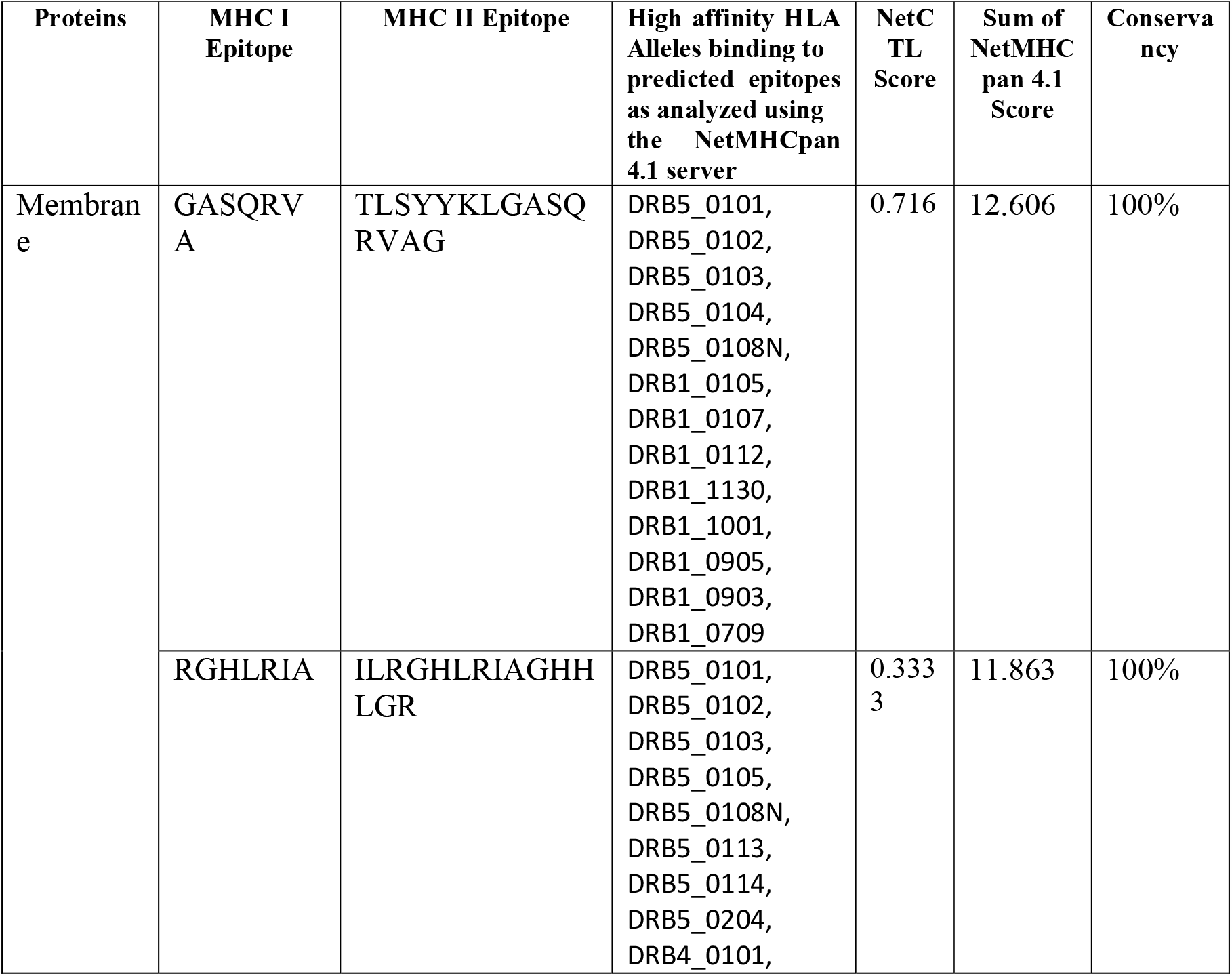
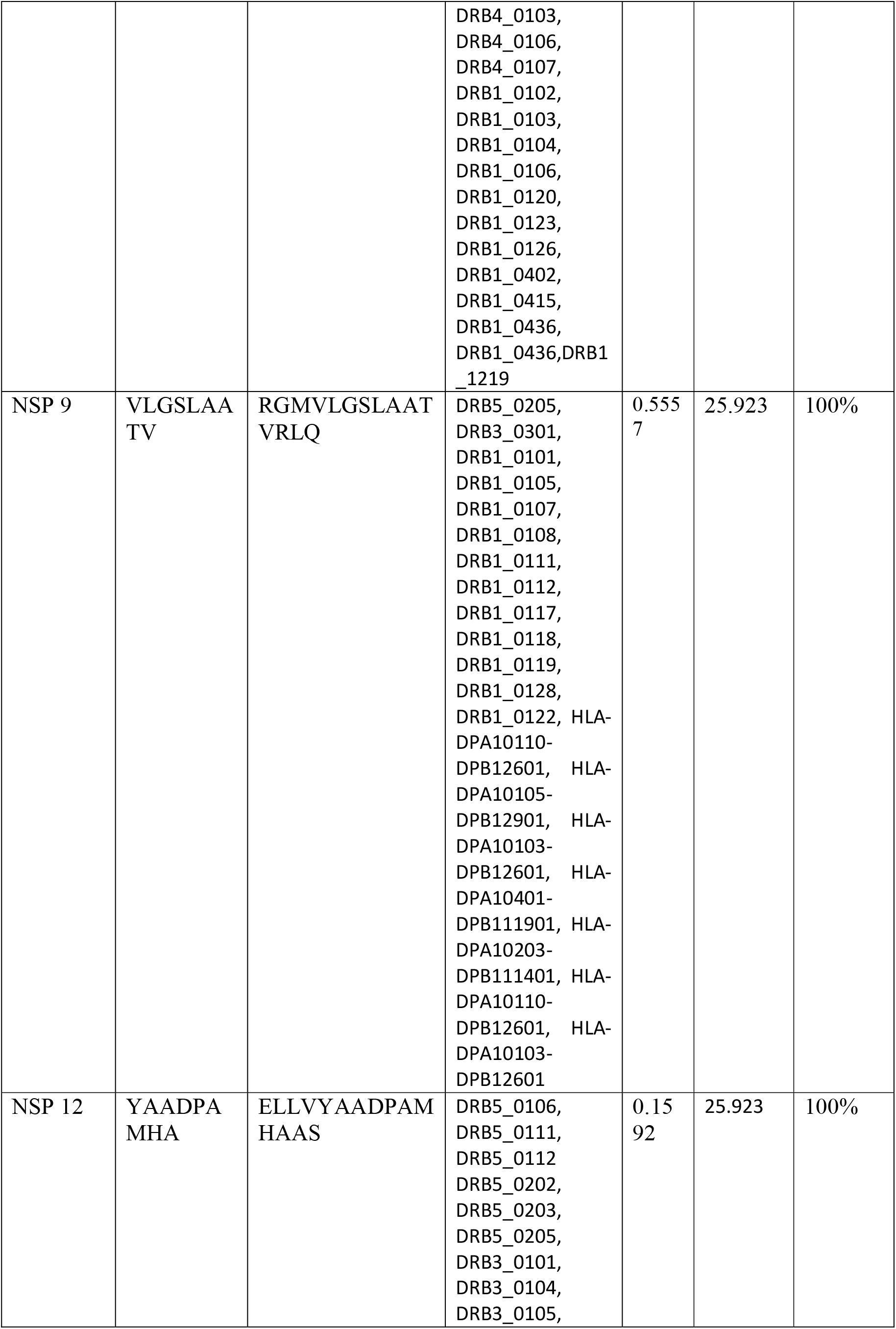
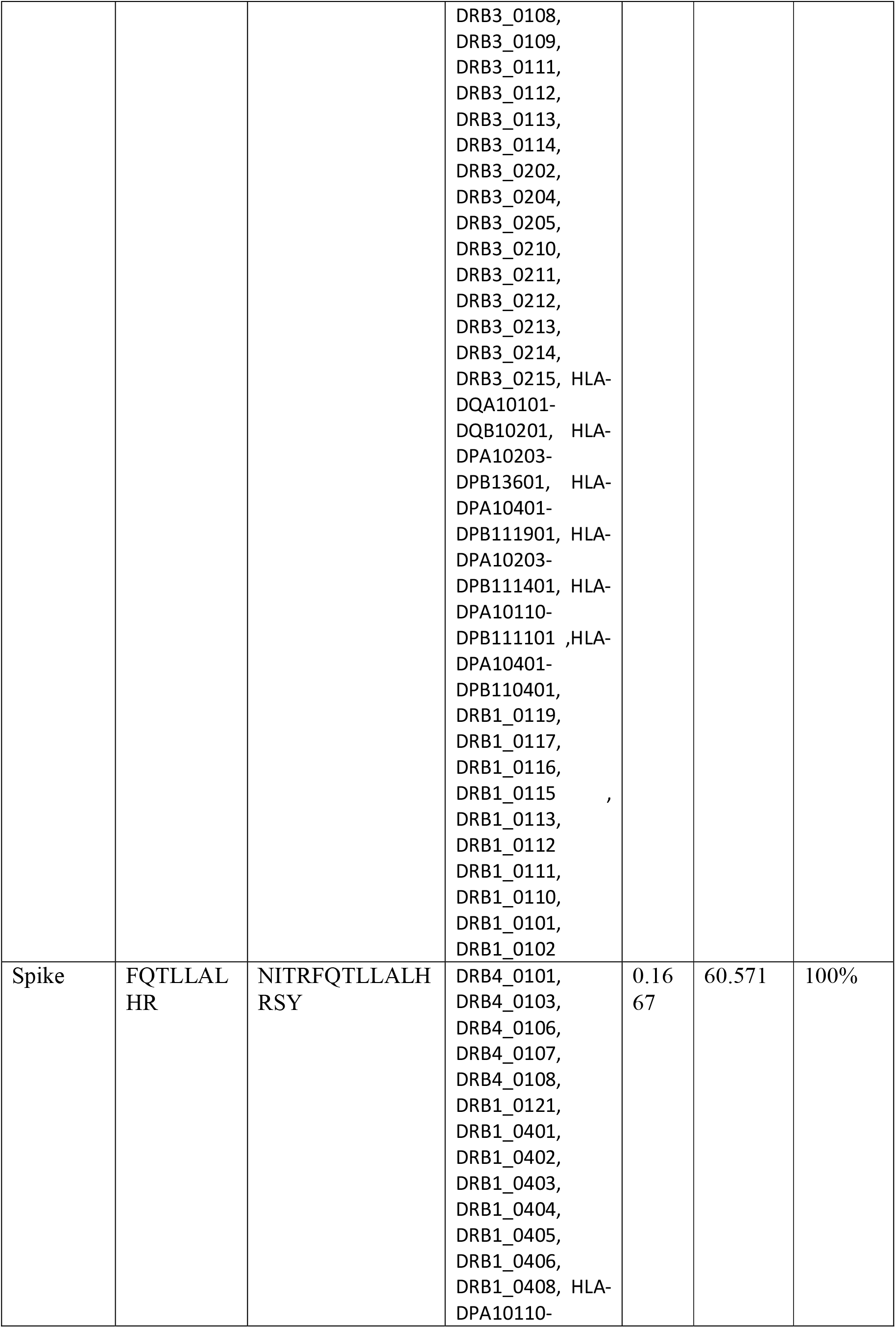
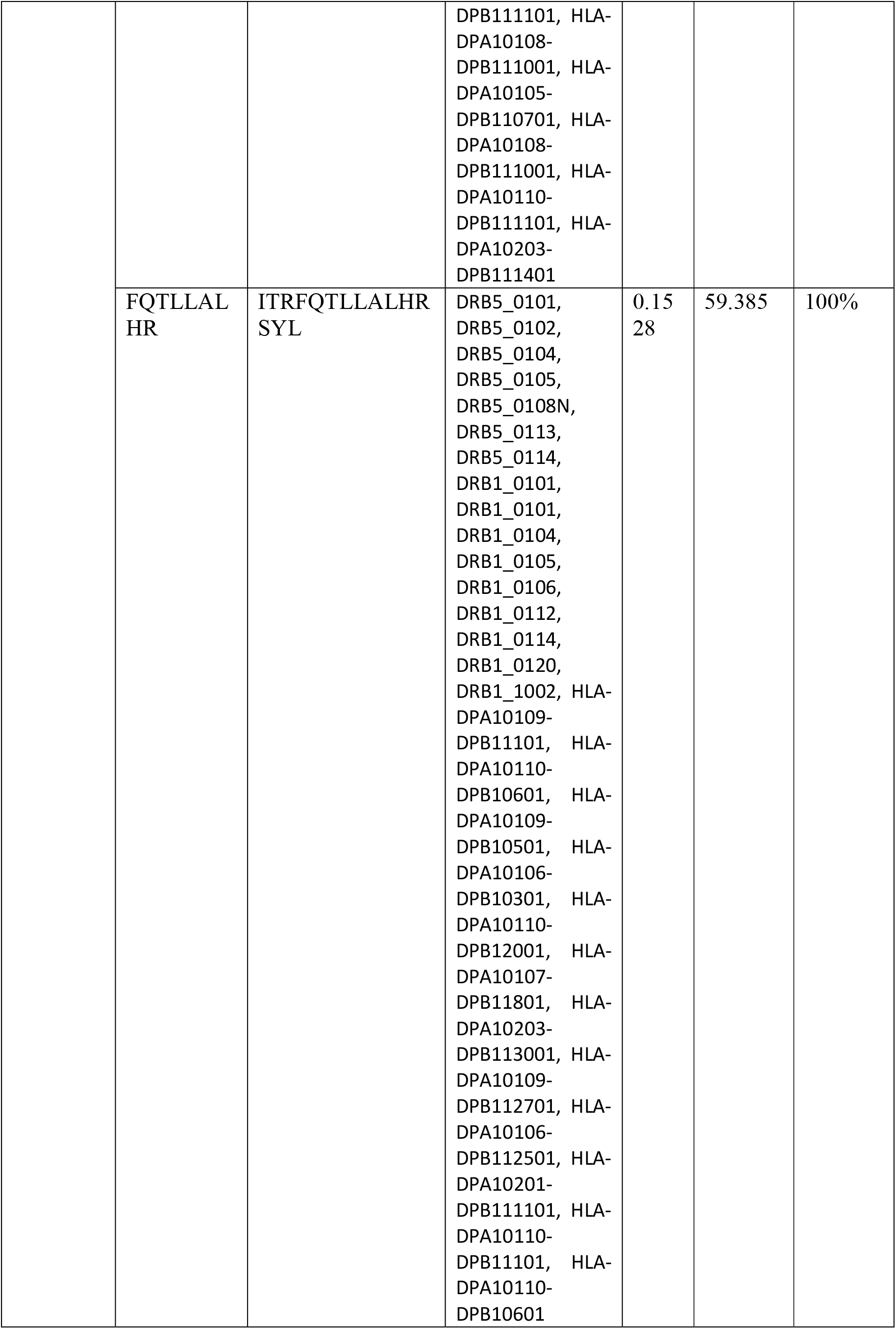
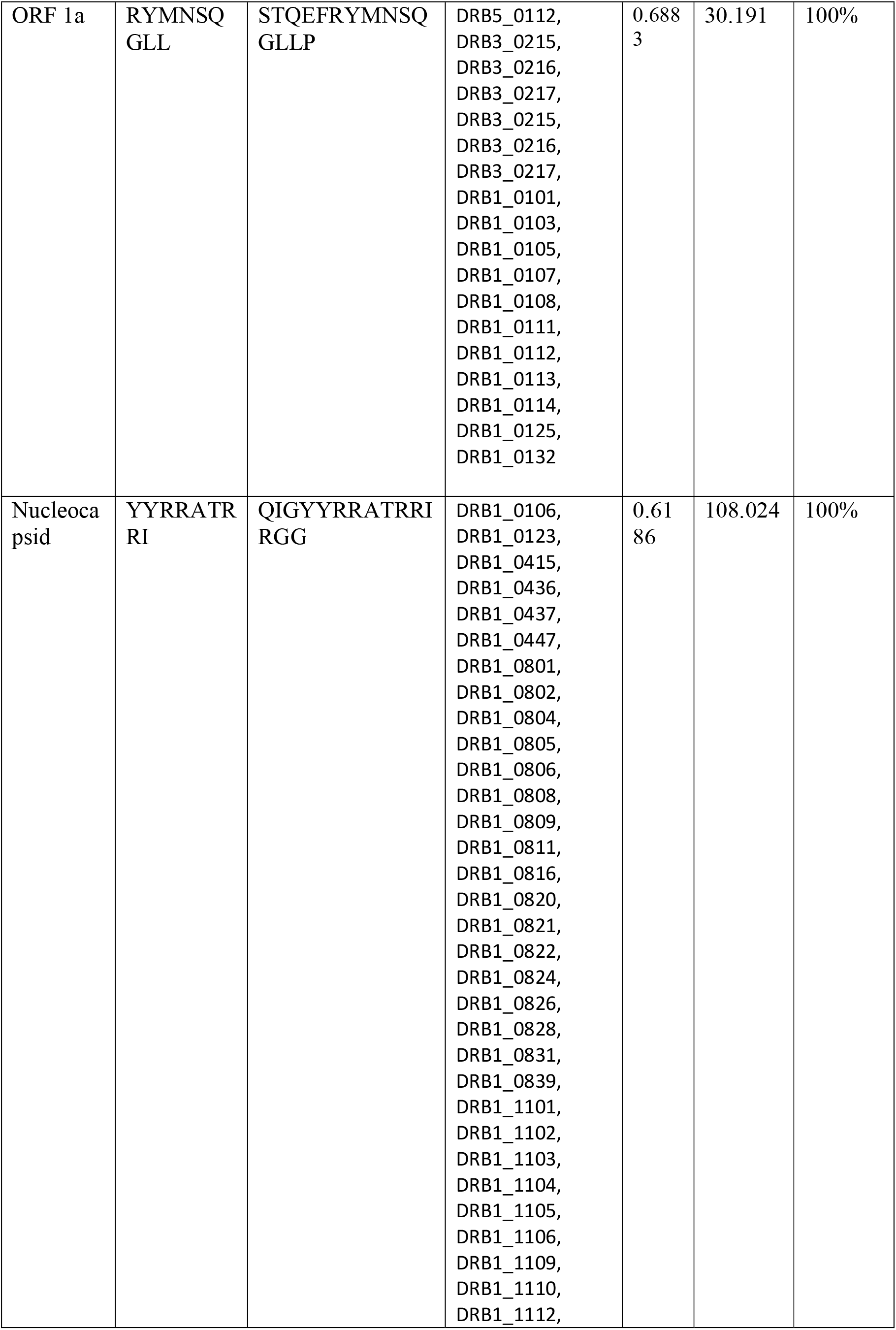
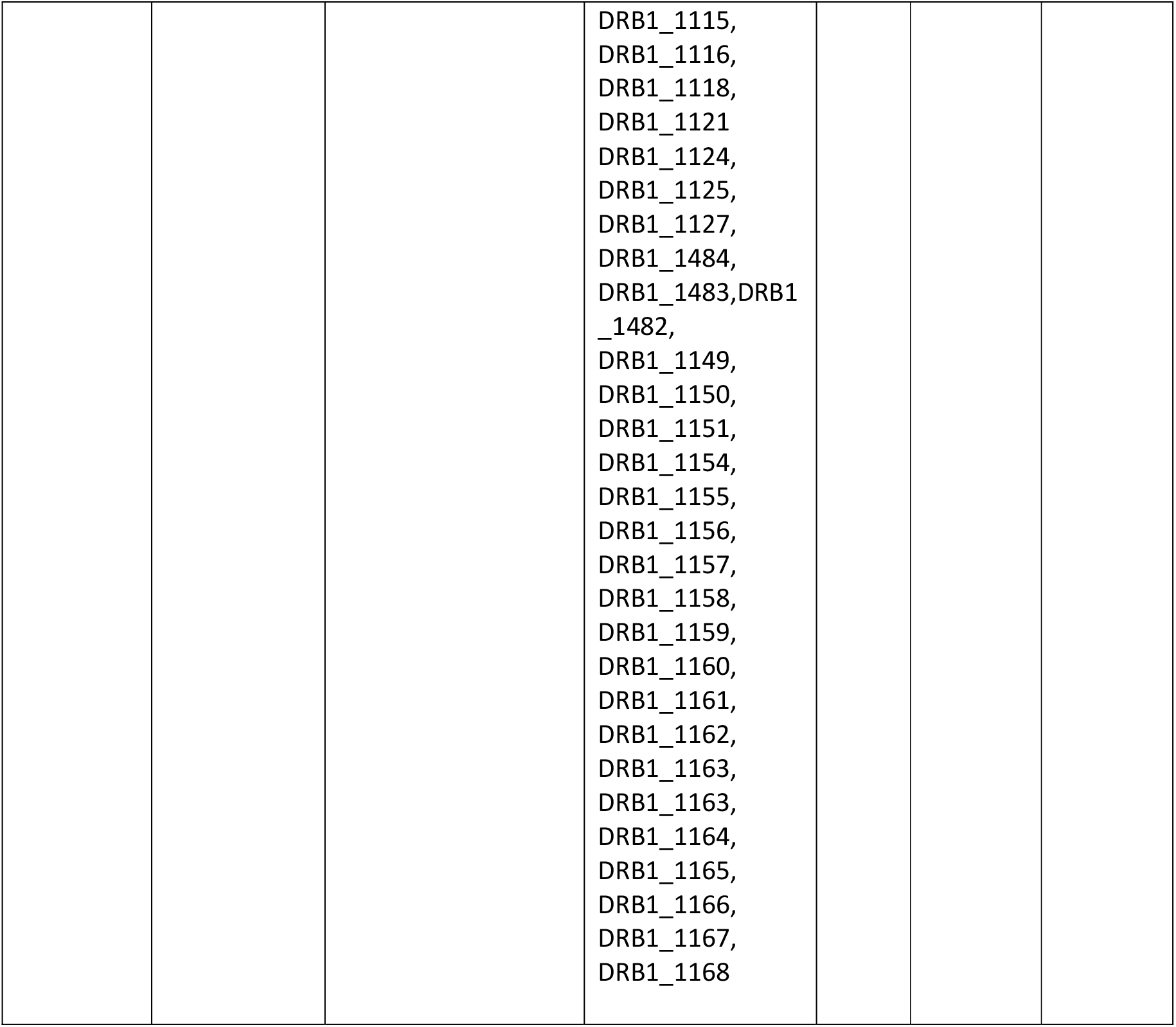
Predicted CTL epitopes from SARS-CoV-2 proteins to design vaccine candidates with their corresponding MHC Class I and II alleles and their immunogenic properties.

### Interferon-© **(IFN-**©**) inducing epitope prediction**

The IFN-© inducing epitopes were predicted from the IFN epitope server. A total of 766 potential MHC II epitopes among which 352 showed positive results. This server executes such prediction with the usage of SVM and MERCI software. Some selected epitope lists were given in Table 4.

**TABLE 4:**
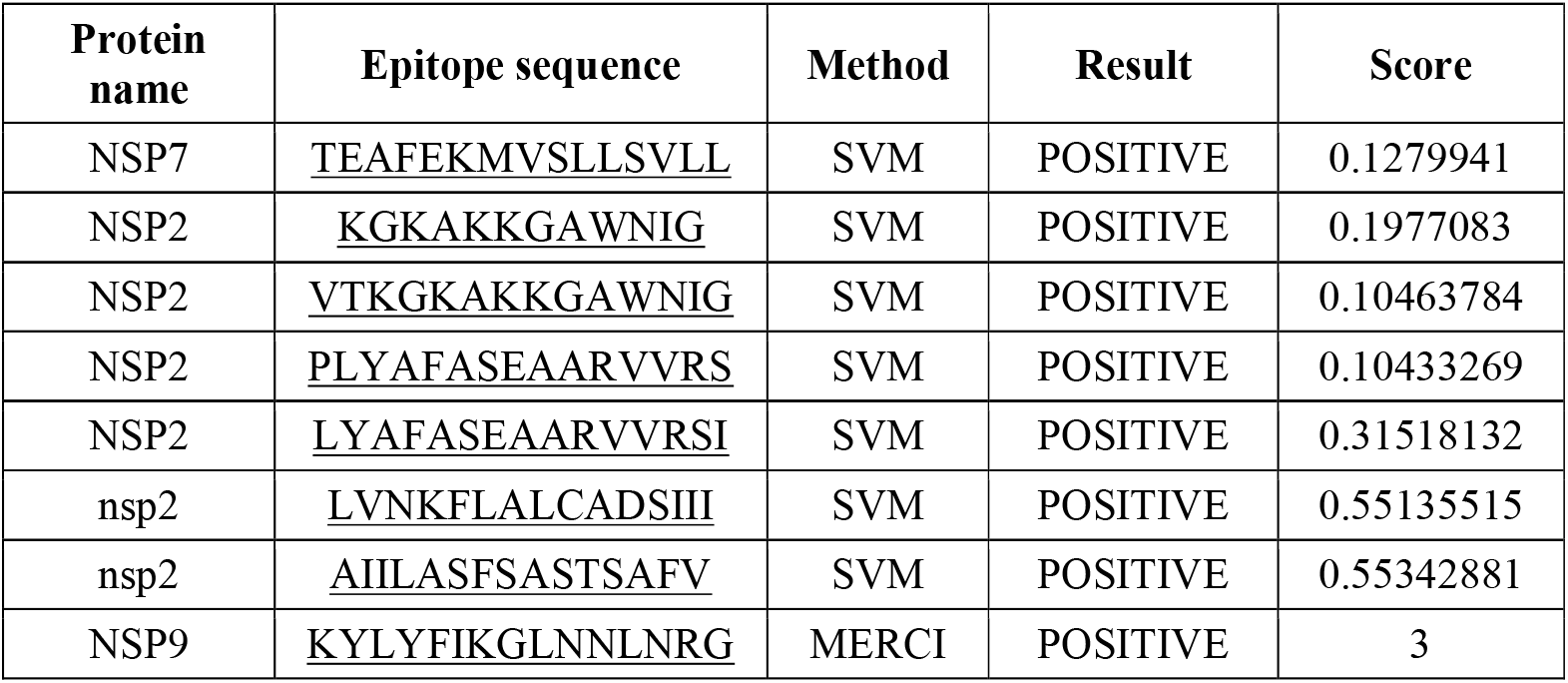
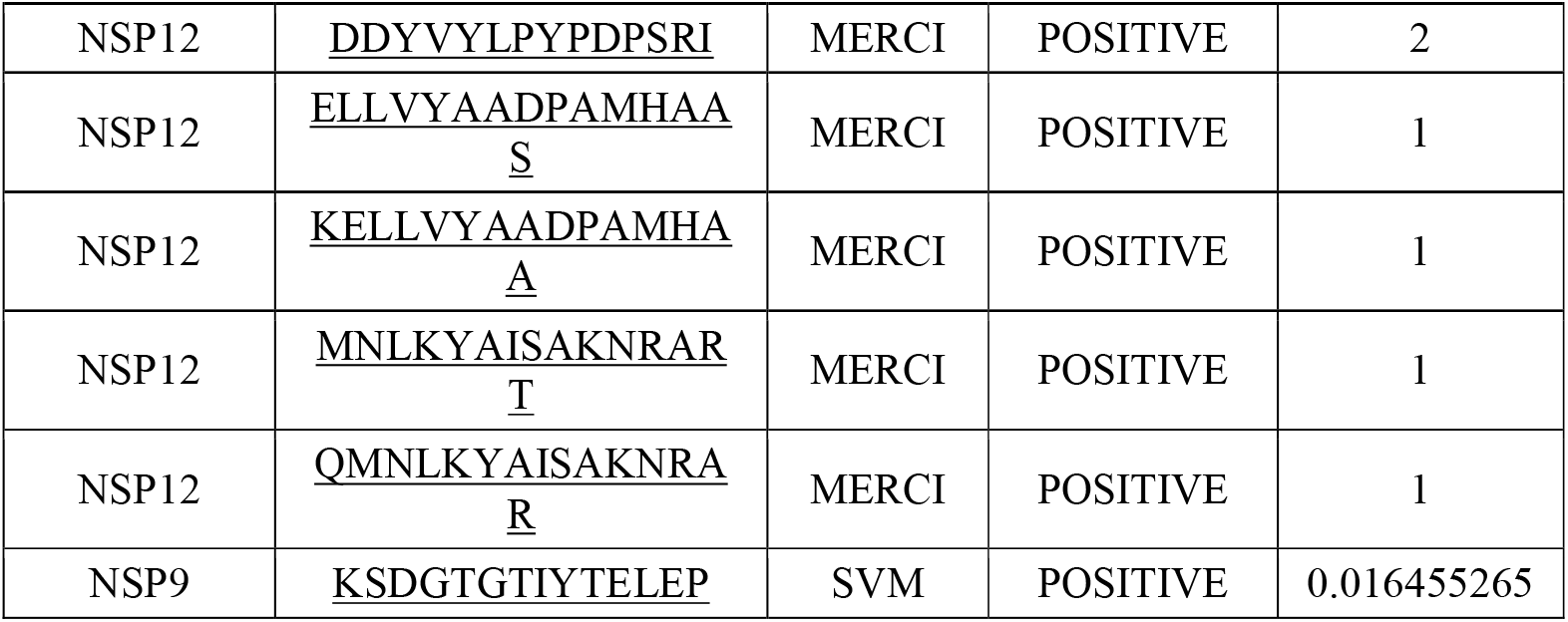
Interferon-© (IFN-©) inducing epitopes.

### Antigenicity, Conservancy and allergenicity assessment

The antigenicity nature of vaccine candidates has a vital role in binding ability with the B and T cell receptor and respective immune response in the host cell. Here we used VaxiJen v2.0 to predict the antigenicity of the screened proteins. And found antigenicity 0.5999 with a 0.4% threshold. The findings indicate the designed protein sequences without adjuvant are antigenic in nature. Similar type of report was reported by Dimitrov et al. where allergen proteins induce an IgE antibody response. We should take care that vaccine candidates should not show any allergic reaction in study. For that allergenicity of the sequence was predicted by the Allertop tool. Our potential protein was not distinguished as an allergen in Allertop based data. All the peptides were subjected to check the conservancy globally, so as to check the conservancy nature of the predicted epitopes was evaluated between all the strains present globally at the IEDB analysis resource by using epitope conservancy analysis tool. Among all 286 peptides showed 100% conserved at a specified identity level are hence taken. We also checked the allergenicity for ensuring the candidate vaccine, which must not stimulate any allergic reactions after introduced into the host body. As predicted by AllerTOP Allergen PHP web servers, as a result we got 148 vaccine candidates was found to be non-allergen (Table S2).

### Modeling of epitopes 3D Structure

The three dimensional structures of all the 148 non-allergenic and 100% conserved epitopes were modeled using PEP-FOLD server and the receptor TLR proteins were modelled using Phyre2 server and further refinement was done(Figure 3).

**FIGURE 3:**
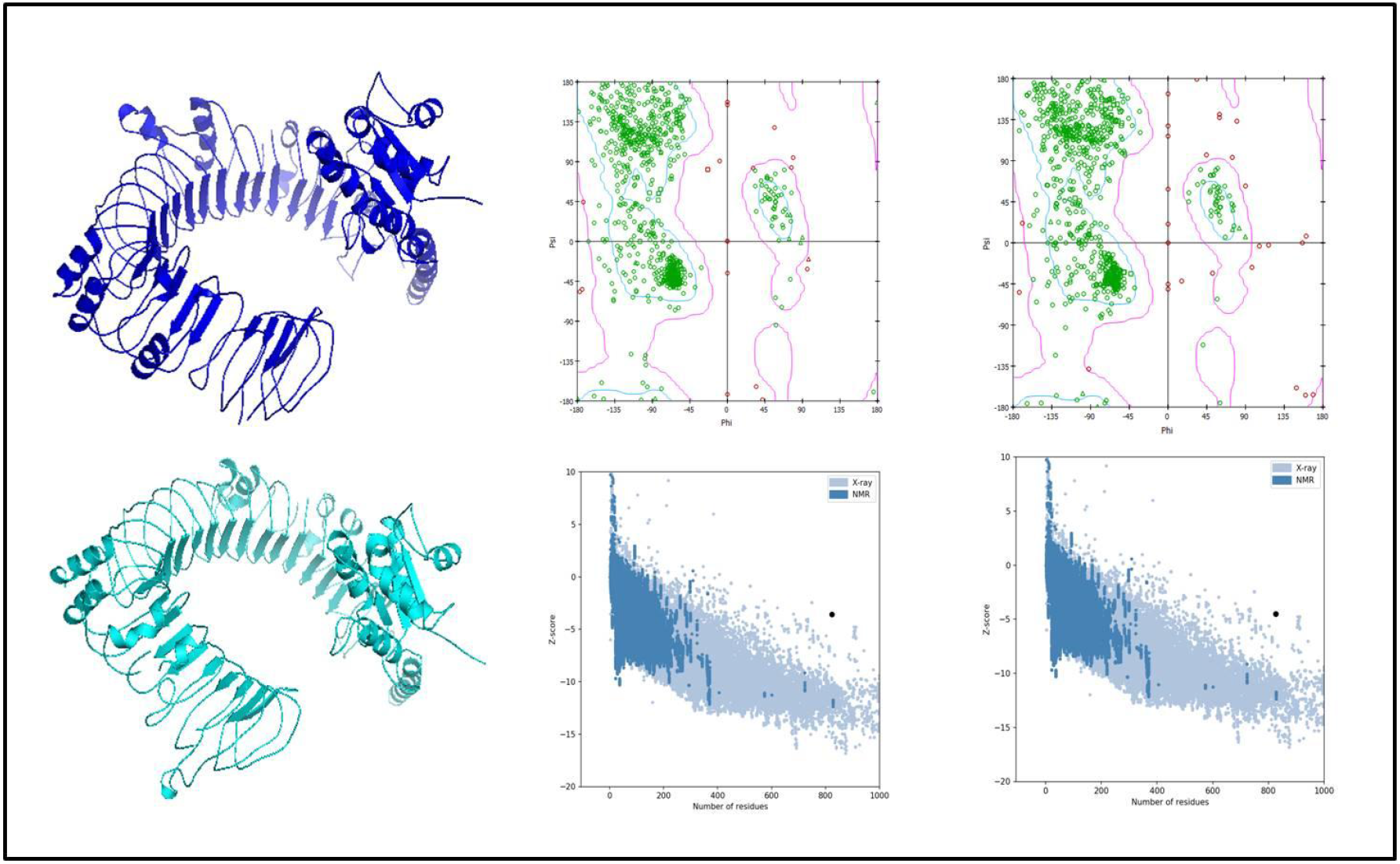
Modeled structure and refinement of human receptor TLR 7 and TLR8. (A) Three dimensional model of TLR 7 AND TLR 8 obtained by homology modeling (B) PROSA analysis of TLR 7 and TLR 8 (C) Ramachandran plot analysis of Receptor proteins showing favored, allowed (4.8%), generously allowed (2.6%), and disallowed (1%) regions.

### Interaction of Predicted Vaccines with Potential human Receptors

The human receptor proteins and the CTL epitopes binding affinity were obtained by Molecular docking. 10 human receptors proteins were docked out of 497 predicted CTL epitope including the furin and all TLRS. Out of 497, only 56 predicted CTL epitopes gave strong binding affinities in terms of attractive van der Waals global energy. Among all, seven epitopes were obtained having elevated degree of conservancy; antigenic amino acid residues and also positive IEDB immunogenicity score (Table 4). These above explained character of an epitope based peptide can be a promising peptide vaccine candidate. Besides, the rest of the epitopes have shown poor binding affinities. All seven epitopes modelled using Pep Fold server was docked with human Toll-like receptors (TLRs). The predicted CTL epitopes “ITRFQTLLALHRSYL” and “RGMVLGSLAATVRLQ” exhibited the highest binding affinity of−6.8 kcal/mol with both TLR8 and TLR2 respectively and “ITRFQTLLALHRSYL” had −6.5 kcal/mol of binding energy with TLR7(Figure 4) while “ELLVYAADPAMHAAS” exhibited the lowest binding affinity of −4.8 kcal/mol with TLR 9(Table 5).The Docking energies of previously mentioned epitopes which is having lowest global energy are nowhere very close to those of top seven epitopes. Moreover, the post-docking result uncovered the presence of five hydrogen bonds in protein-peptide complexes inside a distance of 3.68 Å-5.0 Å, which is bringing up the solidness of the docked complex. Furthermore, the unwavering quality of the docked complex looks to be well preserved with hydrogen bond. in our study we found 7 epitopes that can be taken as potential vaccine candidate for the epitope based vaccine design because of their strong binding affinity within the groove of protein, lowest energy, which point out the strength of docked complexes. Furthermore, this peptide was discovered to be conserved among worldwide strains of SARS-COV-2.

**TABLE 5:**
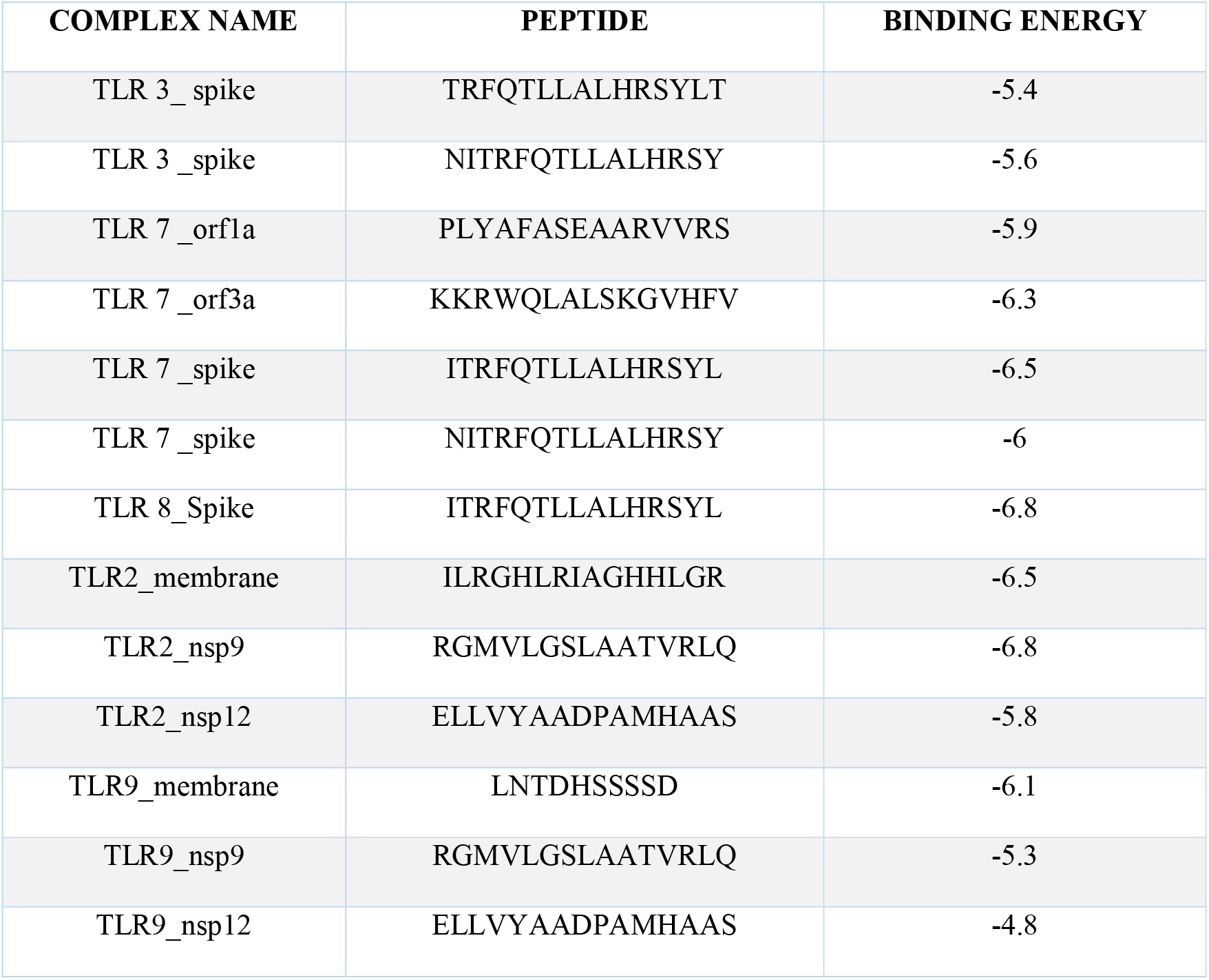
Interaction studies of modeled CTL epitopes with Human TLRs.

**FIGURE 4:**
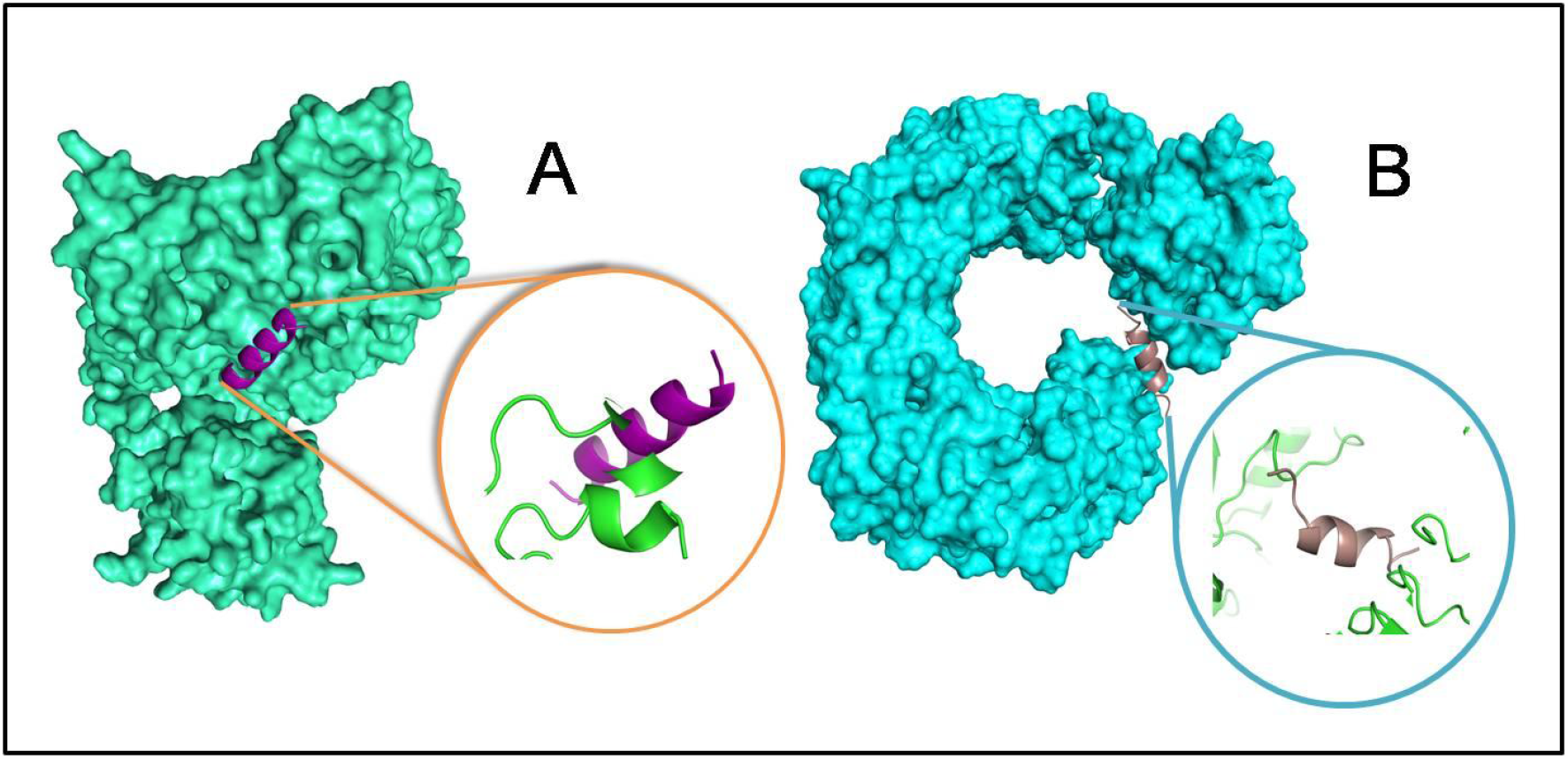
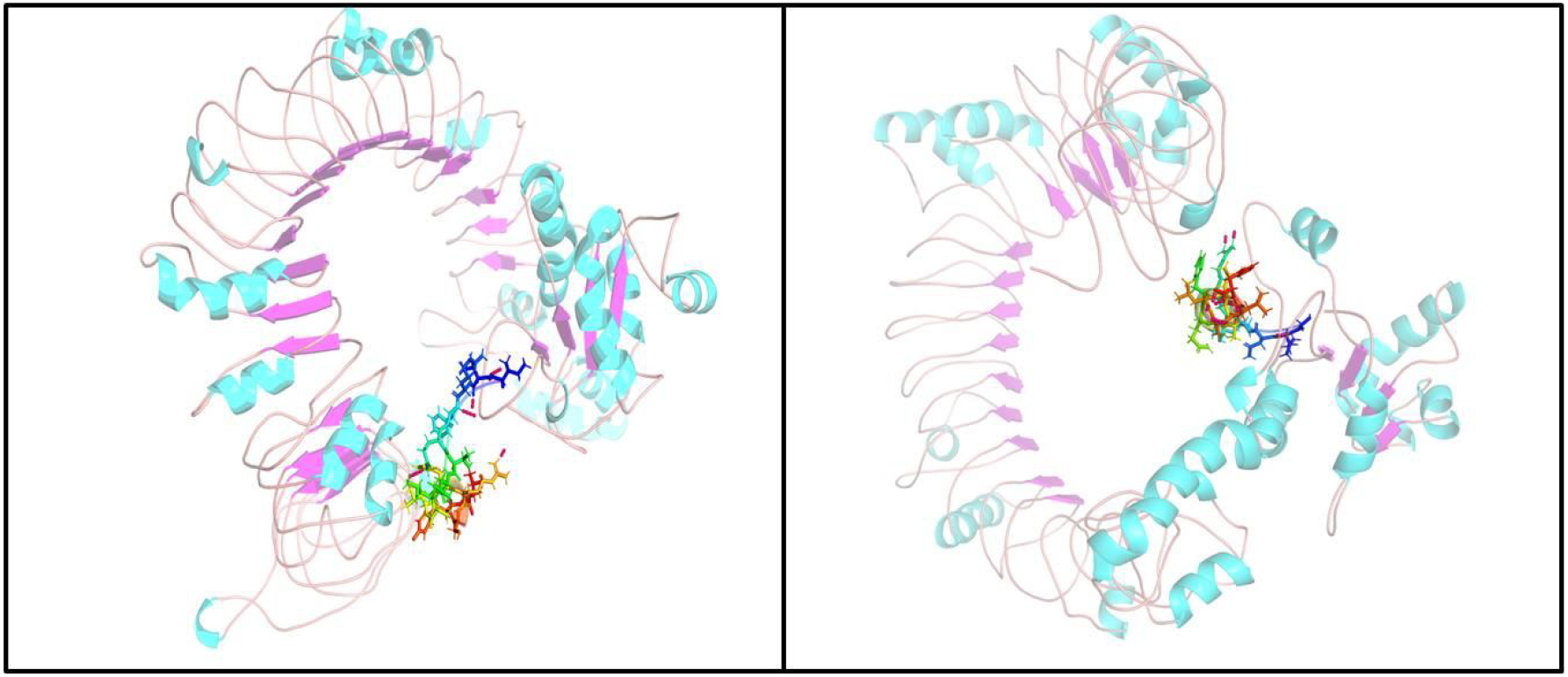
Representative model of interaction between predicted CTL and HTL epitope through molecular docking studies (A) ITRFQTLLALHRSYL with TLR 7, binding energy = −6.5 kcal/mol. (B) ITRFQTLLALHRSYL with TLR 8, binding energy = −6.8 kcal/mol.

### Molecular dynamics simulation

To explore the physical movements of atoms and molecules of the ultimate epitope vaccine candidate, we performed the molecular dynamics simulation. We mainly took care of its density, pressure, temperature, energy components and volume, which were evaluated and their stabilities were confirmed by this dynamic simulation process the stability of the receptor proteins complexes and multiepitope vaccines candidate as ligand was studied through root mean square deviation of the protein backbone and all side chain residues respectively, of the multi-epitope antibody develop for the time-frame of 10 ns. Further to analyze the stability of the ligand receptor interaction the RMSF of amino acid side chains was noted. We found here very mild fluctuations in the RMSD graph, which indicates the uninterrupted interaction between ligand and receptor. In other hand, the RMSF plot showed high fluctuations indicating the flexibility in the regions among ligand-receptor complexes. The RMSF peaks were found to be very elevated, greater than 6.4Å, which showed high degree of flexibility in the receptor-ligand complex. The simulation also monitored the secondary structure elements of proteins(Figure 5).

**FIGURE 5:**
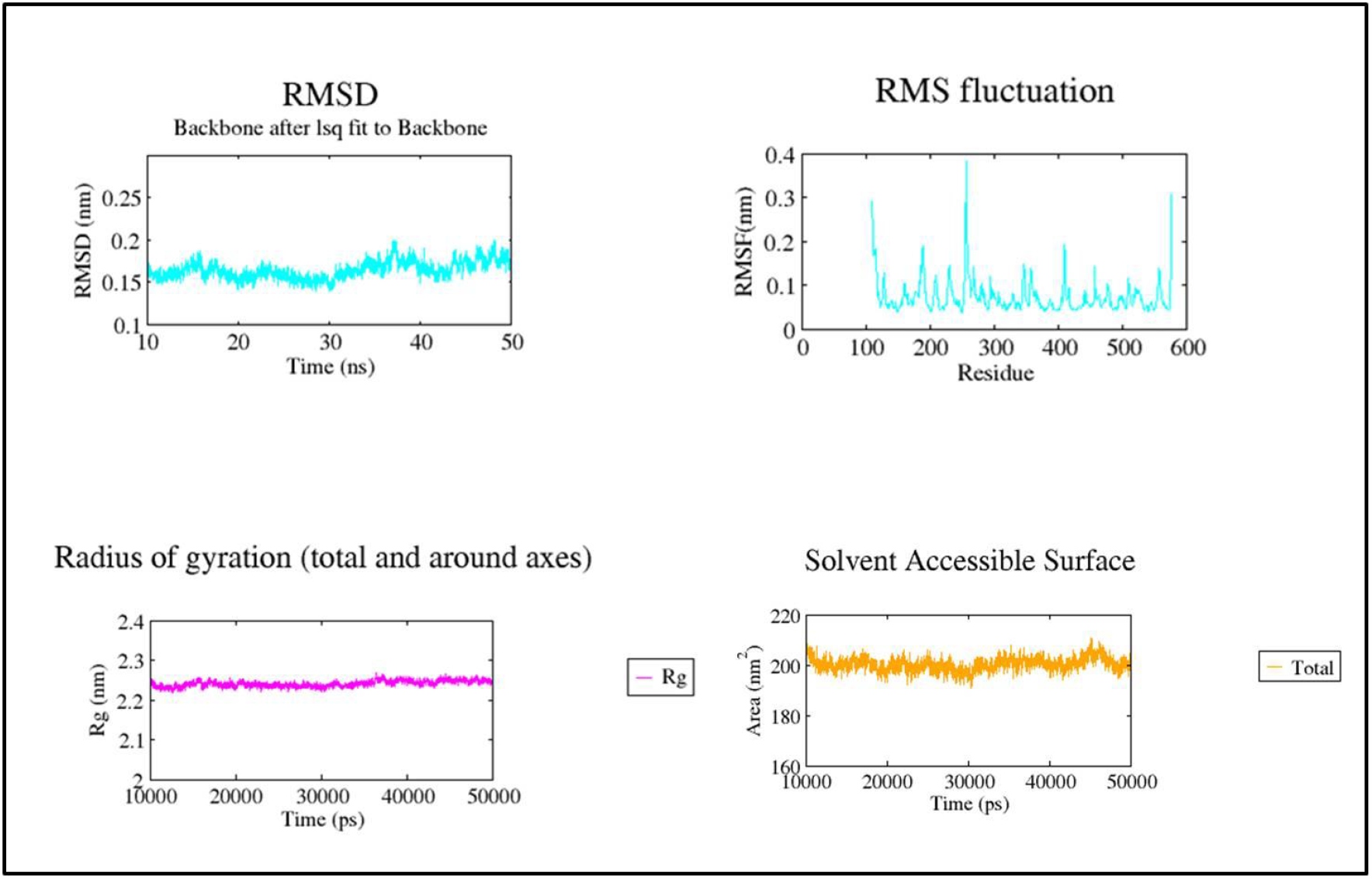
Molecular dynamics simulation study of TLR7, TLR 8 and epitope complex. (A) Root Mean Square Deviation (RMSD) with a time duration of 20 ns, (B) Root mean Square Fluctuation (RMSF) of the docked complex side-chain with a time.

### Population Coverage

The expected population coverage is the percentage of individuals in the population who are likely to induce an immune response to at least one T cell epitope in the set. In order to identify the epitope set associated with the MHC allele that would maximize population coverage, we took a greedy approach: (i) We first identified the MHC allele with the highest individual population range and then the set associated with the epitope was initialized (ii) We gradually added epitopes associated with other MHC alleles, which result in the greatest increase in accumulated population coverage. As a result, all epitopes had a mean combined Class I and Class II coverage of 94%. This step was performed using the entire data set for the world population, and the MHC restricted allele used in this case is (A ∗ 01: 01, A ∗ 02: 01, A ∗ 03: 01, A ∗ 24: 02, A ∗ 26: 01, B ∗ 07: 02, B ∗ 08: 01, B ∗ 27:05, B ∗ 39: 01, B ∗ 40: 01, B ∗ 58: 01, B ∗ 15: 01, HLA -DRB1 ∗ 03: 01, HLA DRB1 ∗ 07: 01 and HLA-DRB1 ∗ 15: 01) (Table 6)(Figure 6A and 6B).

**TABLE 6:**
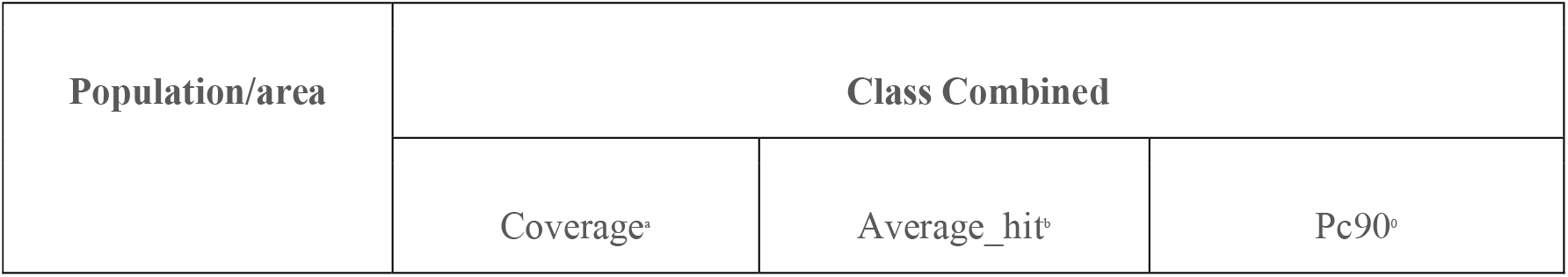
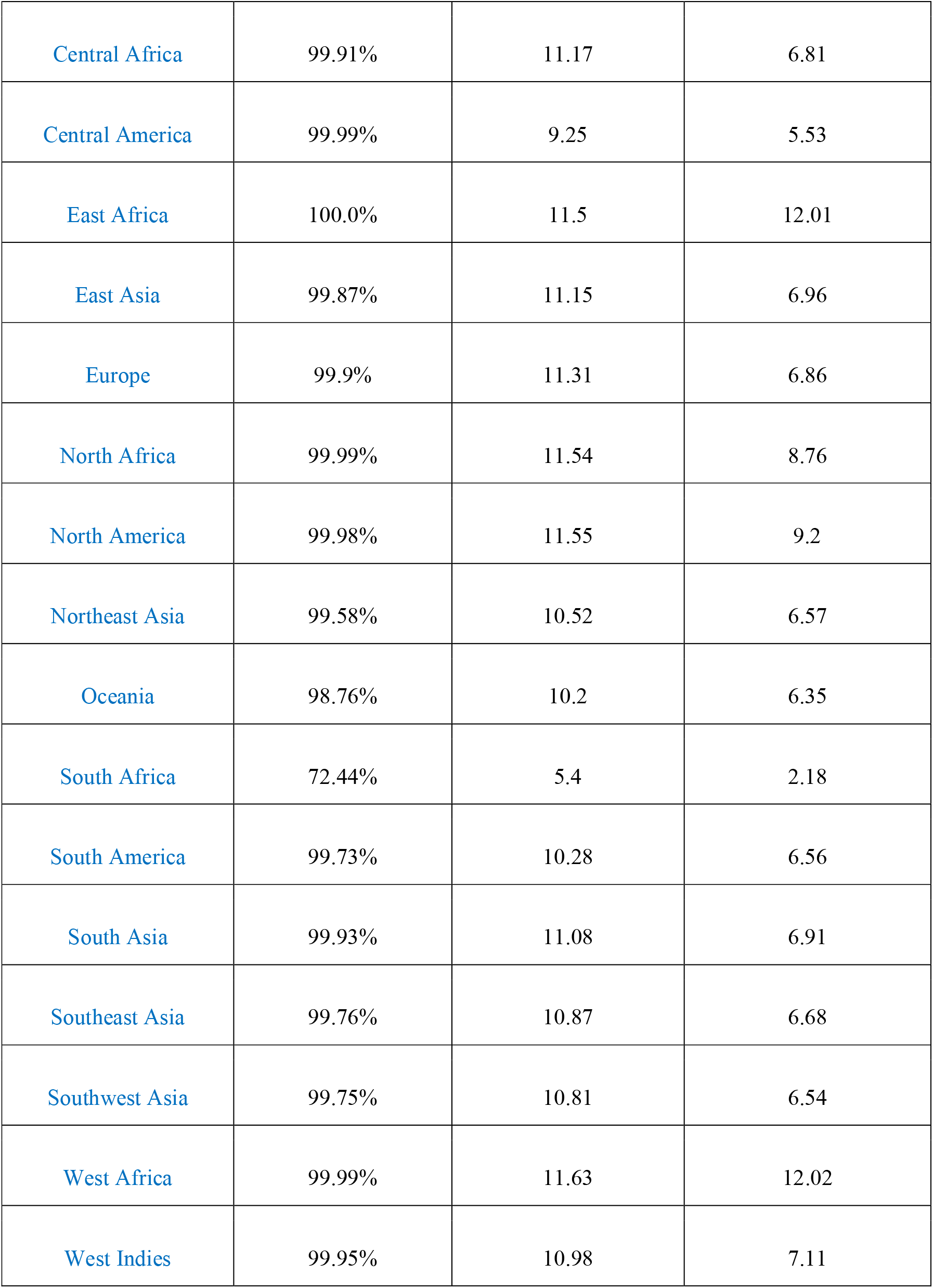
Population coverage of the selected epitopes of the vaccine construct, as predicted by IEDB server.

| Population/area | Class Combined |  |  |
| --- | --- | --- | --- |
|  | Coverage<sup>a</sup> | Average_hit<sup>b</sup> | Pc90<sup>c</sup> |
| Central Africa | 99.91% | 11.17 | 6.81 |
| Central America | 99.99% | 9.25 | 5.53 |
| East Africa | 100.0% | 11.5 | 12.01 |
| East Asia | 99.87% | 11.15 | 6.96 |
| Europe | 99.9% | 11.31 | 6.86 |
| North Africa | 99.99% | 11.54 | 8.76 |
| North America | 99.98% | 11.55 | 9.2 |
| Northeast Asia | 99.58% | 10.52 | 6.57 |
| Oceania | 98.76% | 10.2 | 6.35 |
| South Africa | 72.44% | 5.4 | 2.18 |
| South America | 99.73% | 10.28 | 6.56 |
| South Asia | 99.93% | 11.08 | 6.91 |
| Southeast Asia | 99.76% | 10.87 | 6.68 |
| Southwest Asia | 99.75% | 10.81 | 6.54 |
| West Africa | 99.99% | 11.63 | 12.02 |
| West Indies | 99.95% | 10.98 | 7.11 |

**FIGURE 6A and 6B:**
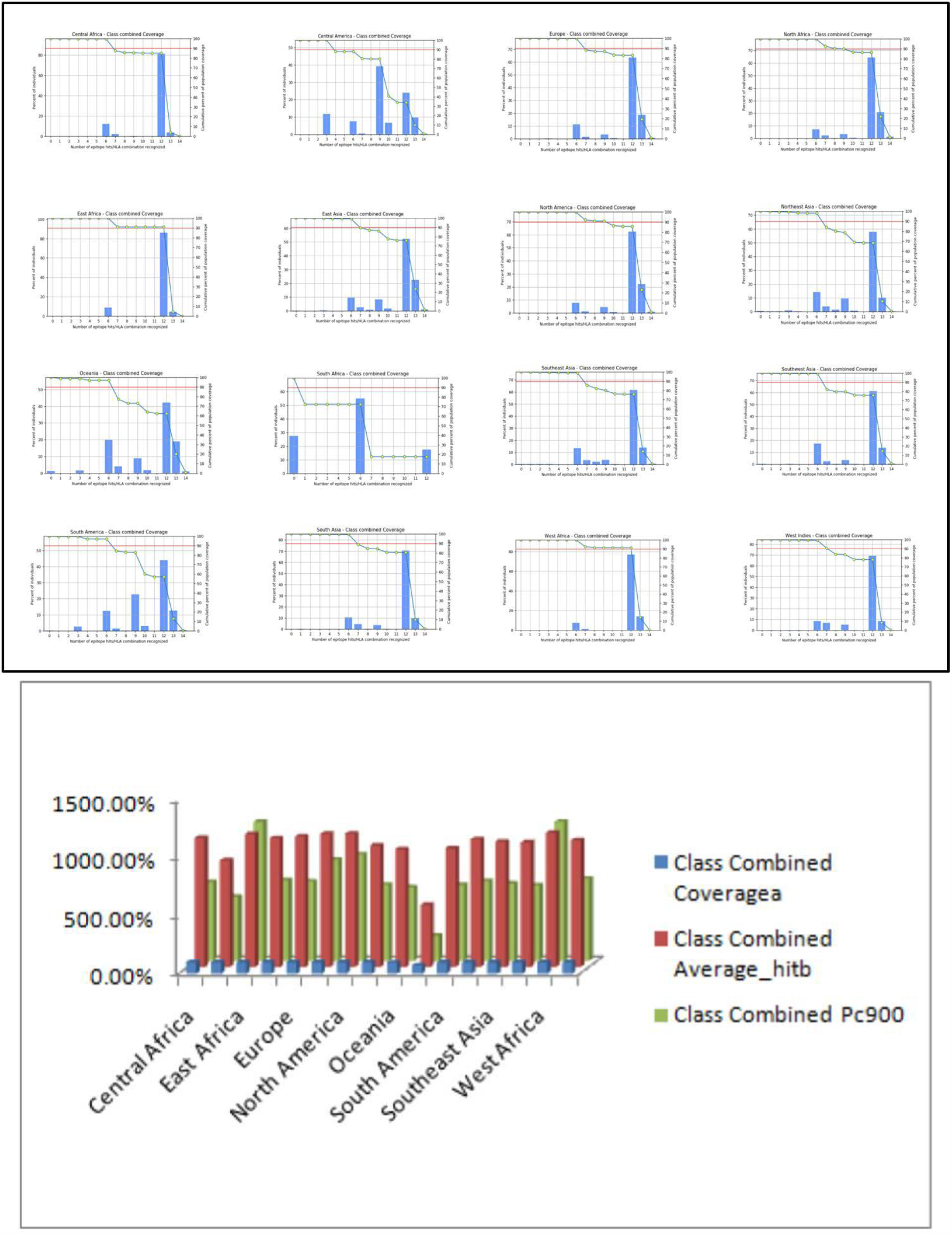
Population coverage of the selected epitopes of the vaccine construct, as predicted by IEDB server.

## DISCUSSION

The purpose of this analysis is to recognize potential crossreactive vaccine candidates and defensible using a holistic approach to immune informatics and reverse vaccinology. The reverse vaccinology approach can be useful to reduce the time frame for vaccine candidate discovery and advantages over traditional methods which tend to Focus more on the development of pathogens and protein extraction, and this large-scale testing of proteins is expensive and time consuming [63, 64, 65].Researchers have identified many in-silico vaccine candidates that can generate enormous clinical and pre-clinical outcomes [66]. In this study, suspected antigenic proteins identified from B cells (linear and conformal) as well as T cell epitopes (HTL and CTL) may possibly build a peptide vaccine for SARS-COV-2 [67, 68]. Our investigation revealed 3,354 proteins as non-redundant proteins, although because of their 60 percent sequencing similarity, 84 proteins were classified as redundant. [69, 70]. The excess protein has a paralogical function which is caused by duplication during evolution. Since replication, abnormalities or mutations in excess protein are less dangerous to the overall health and are of the organism thus not seen as desirable targets for the creation of vaccines [71]. On the other hand, orthological the proteins are not interchangeable and are known to be more conserved in microbial species and strains and may also be suitable vaccine development candidates [72]. In the next step, the non-redundant protein was compared with the human reference protein. It was found that 556 proteins were human homologues, while 2798 proteins were non-human homologues. Removal of homologous host proteins is important because targeting these proteins will elicit a powerful host immune response. In comparison, homologous host proteins can be integrated and recombined to the host genome and are therefore avoided for vaccine production [73]. Identifying essential and non-homologous proteins is very important since most vaccines attack the pathogen’s key cellular processes. The key proteome is a collection of proteins that are required to support cell life and thus have a wider therapeutic potential [74]. Core pathogen proteins were classified using the CD-HIT “multiple alignments clustering approach” study and had 31 protein clusters with 60 percent identity.

Recent developments in immunological bioinformatics have culminated in tools and servers that can decrease the costs and time of producing conventional vaccines. The development of the effective epitope-based vaccines remains difficult due to the trouble in identifying appropriate immunodominant and epitope candidates [75]. Appropriate prediction of the antigenic epitope of the protein target using an immunoinformatics approach is therefore important for the design of the vaccine on an epitope basis [76, 77].

Here we investigate the development of epitope-based vaccines for SARS-CoV-2 structural proteins (spike, membrane, nucleocapsid, non-structural protein, ORF and envelope). This protein plays an important role In the period of replication and the structure of virus particles. The spike Protein plays a significant role In the binding of the virus to the receptors on the host surface and in sequential fusion to induce viral entry into the host cell TLR7 and 8 [78, 79, 80]. Membrane and envelope proteins are essential for replication, particle collection in human cells, and entry of viruses [81, 82]. Corona virus nucleocapsids are structural proteins that form complexes with genomic RNA, interact with viral membrane proteins during virion assembly and play an important role in increasing the efficiency of transcription and viral assembly [83]. NS proteins (NS1 and NS2) formed by a smallest segment of RNA present in virus-infected cells. NSP is transported to the nucleus and involved in the Inhibiting host mRNA synthesis, core viral mRNA export, and viral protein translation, but is not found in viral particles. NSPs play an important role in viral virulence by combatting the work of a host interferon. Five major open reading frames (ORF) are encoded by the coronavirus genome, including a 5 ‘shifted polyprotein skeleton (ORF1a/ORF1ab) and four 3’ canonical structural proteins essential to all spike(S), membrane(M), envelope(E), and nucleocapsid(N) proteins, including all coronaviruses. The 3D models of the construction were then created and an online server was used and then tested by a Ramachandran plot analysis. Moreover, with the epitope vaccine docking research various toll-like receptors such as TLR2, 3 7, 8 & 9 were performed. We aim TLR7, and mostly TLR8. In the entire antiviral reaction, both TLR7 and 8 are significant participants. Dendritic plasmacytoid cells (pDCs) and B cells are activated by TLR7-specific agonists and primarily induce IFN-and IFN-regulated cytokines.TLR8-specific agonists activate myeloid DCs, monocytes, even monocyte-derived DCs, and primarily lead to the release of proinflammatory cytokines and chemokines. The study was performed on the structural protein levels of primary, secondary and tertiary. Conserved B cell epitopes were forecast by IEDB research resource and BepiPred 2.0 [48]. The epitope location in the 3D structure of the protein was visualized by Pymol [59].They were tested to further improve the allergenic and physicochemical properties of the expected epitopes. Analysis of digestion indicates that the peptides administered were stable and safe to use during the analysis.

A good vaccine should have strong physicochemical properties in its development, formulation, storage and use, in addition to enhanced immune response capability. Formational B cell epitopes play a crucial role in induction humoral reactions. The high propensity of this structure to activate B lymphocytes is shown by the presence of a large number of B cell epitopes within the vaccine molecule. In order to successfully pass the vaccine protein to the antigen-presenting cells, the binding affinity of the vaccine to the immune receptor (TLR7 and 8) is important. The results of the docking analysis revealed a strong interaction between the vaccine protein and TLR7 and 8.

The outcomes of the antigenicity analysis observed vary, which is known to be the essential antigenic capacity of a potent peptide, and all immunoinformatics tests have shown similar numbers. Moreover, the binding domain of HLA-B was found in both experiments to be preserved and reconciled with current research efforts [84, 85].

Potential of the CTL epitopes for SARS-CoV-2 structural proteins have been predicted. For the selected peptides, molecular docking methods were used to evaluate MHC-1 and binding affinities of peptide [86]. The binding affinity of peptide-MHC-I complexes has also been validated by other results, including C-terminal cleavage affinities. 140 peptides have been identified as potential targets for responsive MHC-I protein interactions (HLA-B) with maximum binding affinities and antigenicity. This increases the likelihood of promising targets for potential residue vaccine targets. SARS-CoV-2 structural proteins were measured and cross tested with the IEDB server in terms of surface usability, hydrophobic, surface stability, and antigenicity [87]. Extensive literature studies were conducted and selected peptides against SARS-CoV-2 were not identified. Predicted peptides have been modeled by I-TASSER, ORION: Web Server and PEP-FOLD3 server and MHC-1 server for more refinement using Autodock Vina and Haddock. [88]. PyMOL has been used to check the interactions of docked complexes. In order to test the stability of the peptide receptor complex, a molecular dynamic simulation of the docking complexes (peptides and TLR7 and 8) was performed using the GROMACS 5.1.1 version. Various analyzes were then carried out such as energy minimisation, pressure calculation, temperature measurement and potential energy measurement. During the production phase, the temperature and pressure of MD simulation system were roughly 300K and 1 ambient, using high voltage pulses (duration 10 ns and 1000 ps, amplitude 15 kV, repeat frequency 1–30 kHz) a stable simulation system and a good md run, respectively. MD simulation system temperature and pressure (peptide and TLR-2, 3, 7, 8 and 9 complexes) during production were approximately 10ns and 1000 ps, respectively, implying stable systems and strong MD activity [89]. The complex of root mean square deviation (RMSD) plot represents the structural variations in the overall structure of the peptide complex and the receptor. We have also regularly incorporated epitopes associated with other MHC alleles, which has resulted in the largest increase in total population coverage. As a result, all epitopes had a combined Class 1 and Class 2 coverage of 100 per cent of the world population dataset detected in East Africa followed by Central America, North America and West Africa. Estimates of population coverage are based on binding impacts. for MHC and/or T cell limitation, but the technique can be used more widely. The algorithm has been introduced as a web application that can be used by vaccinations or diagnoses dependent on epitopes can be formulated to maximize population spread, thus attenuating variability (i.e. the number of distinct epitopes included in the diagnosis or vaccine) and minimizing disparities in coverage among different ethnic groups.

The development of a potent vaccine includes a detailed analysis and investigation of immunological associations with SARS-CoV-2 [90, 91]. However, due to the severity and urgency of the COVID-19 outbreak, innovative approaches would not be adequate to serve the urgency of the situation. However in the case of silicon and numerical forecasts, it is helpful to lead researchers in discovering a potential vaccine and helping to monitor COVID-19. Vaccine production is a costly and lengthy process with a high risk of failure and a number of years are needed to produce a successful commercial vaccine. Computational analyzes indicate that the registered epitope-based peptide vaccine could have the ability to protect against SARS-CoV-2 infection.

### Conclusion

TLR has diverse pleiotropic effects on the immune system and has been shown to promote antigen-specific cellular and humoral immunity, TLR7/8 induce the development of IgG and the recruitment of B-cell cytokines and the secretion of **Interferon-**© **(IFN-**©) by natural killer cells. In our research, immuno-informatics techniques were used to design a possible SARS-CoV-2 vaccine candidate consisting of CTL, HTL and **Interferon-**© **(IFN-**©)epitopes that could cause robust immune responses. Possible vaccine candidates for structural, non-structural and accessory SARS-CoV-2 proteins were shown to be both antigenic and immunogenic. The association analysis between TLR receptor and CTL, HTL epitopes, resulted in a good binding affinity of spike protein peptides to TLR 7 and 8. The stability of the vaccine candidate has been confirmed by molecular docking tests. According to many studies, the creation of an epitope-based vaccine is a successful technique that can contribute to the production of vaccines with the potential to cause the necessary immunogenicity. Based on many in vitro and in vivo studies, if scientifically and clinically formulated, epitope-based vaccines provide several advantages over other vaccine types, including quick and accurate design, time-and cost-effective formulations, and ideal immunogenicity with reduced adverse effects. However the findings of this analysis in silicon must be checked using laboratory and animal models.

## Acknowledgement

The authors are thankful to IMS & SUM Hospital, BRAF Help Desk of C-DAC, Pune, for providing a fruitful research facility to carry out the current research work. The AN, MS, AG and BR are also grateful to Siksha ‘O’ Anusandhan (Deemed to be University), Bhubaneswar, Odisha, India, for providing PhD fellowship and all assistance to carry out the research activities.

## Author Contribution

JD conceived the idea and supervised; AN, AG and MS contributed equally and did proteomic analysis and immunoinformatics analysis; BR segregated the primary data; SBB and JNM looked into the resource mobilization to undertake the research; All authors agreed to the manuscript presented.

