## Supplementary material for "Inferring Toll-Like Receptor induced epitope subunit vaccine candidate against SARS-CoV-2: A Reverse Vaccinology approach": Table s1

**TABLE S1: SARS-CoV-2 proteins with their protein ID, Name of protein and Protein sequence in FASTA format**

| **Protein ID** | **Protein name** | **FASTA Sequence** |
| --- | --- | --- |
| **BCB97907.1** | ORF8 protein | >BCB97907.1 ORF8 protein [Severe acute respiratory syndrome coronavirus 2]  MKFLVFLGIITTVAAFHQECSLQSCTQHQPYVVDDPCPIHFYSKWYIRVGARKSAPLIELCVDEAGSKSP  IQYIDIGNYTVSCLPFTINCQEPKLGSLVVRCSFYEDFLEYHDVRVVLDFI |
| **sp\|P0DTC3\|** | ORF 3a | >sp\|P0DTC3.1\|AP3A_SARS2 RecName: Full=ORF3a protein; Short=ORF3a; AltName: Full=Accessory protein 3a; AltName: Full=Protein 3a; AltName: Full=Protein U274; AltName: Full=Protein X1 MDLFMRIFTIGTVTLKQGEIKDATPSDFVRATATIPIQASLPFGWLIVGVALLAVFQSASKIITLKKRWQ  LALSKGVHFVCNLLLLFVTVYSHLLLVAAGLEAPFLYLYALVYFLQSINFVRIIMRLWLCWKCRSKNPLL  YDANYFLCWHTNCYDYCIPYNSVTSSIVITSGDGTTSPISEHDYQIGGYTEKWESGVKDCVVLHSYFTSD  YYQLYSTQLSTDTGVEHVTFFIYNKIVDEPEEHVQIHTIDGSSGVVNPVMEPIYDEPTTTTSVPL |
| **sp\|P0DTC7\|** | ORF 7a | >sp\|P0DTC7.1\|NS7A_SARS2 RecName: Full=ORF7a protein; Short=ORF7a; AltName: Full=Accessory protein 7a; AltName: Full=Protein U122; AltName: Full=Protein X4; Flags: Precursor  MKIILFLALITLATCELYHYQECVRGTTVLLKEPCSSGTYEGNSPFHPLADNKFALTCFSTQFAFACPDG  VKHVYQLRARSVSPKLFIRQEEVQELYSPIFLIVAAIVFITLCFTLKRKTE |
| **sp\|P0DTD2\|** | ORF 9b | >sp\|P0DTD2.1\|ORF9B_SARS2 RecName: Full=ORF9b protein; Short=ORF9b; AltName: Full=Accessory protein 9b; AltName: Full=ORF-9b; AltName: Full=Protein 9b  MDPKISEMHPALRLVDPQIQLAVTRMENAVGRDQNNVGPKVYPIILRLGSPLSLNMARKTLNSLEDKAFQ  LTPIAVQMTKLATTEELPDEFVVVTVK |
| **sp\|P59633** | ORF 3b | >sp\|P59633.1\|NS3B_SARS RecName: Full=ORF3b protein; AltName: Full=Accessory protein 3b; AltName: Full=Non-structural protein 3b; Short=ns3b; AltName: Full=Protein X2  MMPTTLFAGTHITMTTVYHITVSQIQLSLLKVTAFQHQNSKKTTKLVVILRIGTQVLKTMSLYMAISPKF  TTSLSLHKLLQTLVLKMLHSSSLTSLLKTHRMCKYTQSTALQELLIQQWIQFMMSRRRLLACLCKHKKVS  TNLCTHSFRKKQVR |
| **YP_009725255.1** | ORF 10 | >YP_009725255.1 ORF10 protein [Severe acute respiratory syndrome coronavirus 2]  MGYINVFAFPFTIYSLLLCRMNSRNYIAQVDVVNFNLT |
| **sp\|P0DTD3** | ORF 14 | >sp\|P0DTD3.1\|Y14_SARS2  MLQSCYNFLKEQHCQKASTQKGAEAAVKPLLVPHHVVATVQEIQLQAAVGELLLLEWLAMAVMLLLLCCC  LTD |
| **sp\|P0DTC1** | ORF 1a | >sp\|P0DTC1.1\|R1A_SARS2  MESLVPGFNEKTHVQLSLPVLQVRDVLVRGFGDSVEEVLSEARQHLKDGTCGLVEVEKGVLPQLEQPYVF  IKRSDARTAPHGHVMVELVAELEGIQYGRSGETLGVLVPHVGEIPVAYRKVLLRKNGNKGAGGHSYGADL  KSFDLGDELGTDPYEDFQENWNTKHSSGVTRELMRELNGGAYTRYVDNNFCGPDGYPLECIKDLLARAGK  ASCTLSEQLDFIDTKRGVYCCREHEHEIAWYTERSEKSYELQTPFEIKLAKKFDTFNGECPNFVFPLNSI  IKTIQPRVEKKKLDGFMGRIRSVYPVASPNECNQMCLSTLMKCDHCGETSWQTGDFVKATCEFCGTENLT  KEGATTCGYLPQNAVVKIYCPACHNSEVGPEHSLAEYHNESGLKTILRKGGRTIAFGGCVFSYVGCHNKC  AYWVPRASANIGCNHTGVVGEGSEGLNDNLLEILQKEKVNINIVGDFKLNEEIAIILASFSASTSAFVET  VKGLDYKAFKQIVESCGNFKVTKGKAKKGAWNIGEQKSILSPLYAFASEAARVVRSIFSRTLETAQNSVR  VLQKAAITILDGISQYSLRLIDAMMFTSDLATNNLVVMAYITGGVVQLTSQWLTNIFGTVYEKLKPVLDW  LEEKFKEGVEFLRDGWEIVKFISTCACEIVGGQIVTCAKEIKESVQTFFKLVNKFLALCADSIIIGGAKL  KALNLGETFVTHSKGLYRKCVKSREETGLLMPLKAPKEIIFLEGETLPTEVLTEEVVLKTGDLQPLEQPT  SEAVEAPLVGTPVCINGLMLLEIKDTEKYCALAPNMMVTNNTFTLKGGAPTKVTFGDDTVIEVQGYKSVN  ITFELDERIDKVLNEKCSAYTVELGTEVNEFACVVADAVIKTLQPVSELLTPLGIDLDEWSMATYYLFDE  SGEFKLASHMYCSFYPPDEDEEEGDCEEEEFEPSTQYEYGTEDDYQGKPLEFGATSAALQPEEEQEEDWL  DDDSQQTVGQQDGSEDNQTTTIQTIVEVQPQLEMELTPVVQTIEVNSFSGYLKLTDNVYIKNADIVEEAK  KVKPTVVVNAANVYLKHGGGVAGALNKATNNAMQVESDDYIATNGPLKVGGSCVLSGHNLAKHCLHVVGP  NVNKGEDIQLLKSAYENFNQHEVLLAPLLSAGIFGADPIHSLRVCVDTVRTNVYLAVFDKNLYDKLVSSF  LEMKSEKQVEQKIAEIPKEEVKPFITESKPSVEQRKQDDKKIKACVEEVTTTLEETKFLTENLLLYIDIN  GNLHPDSATLVSDIDITFLKKDAPYIVGDVVQEGVLTAVVIPTKKAGGTTEMLAKALRKVPTDNYITTYP  GQGLNGYTVEEAKTVLKKCKSAFYILPSIISNEKQEILGTVSWNLREMLAHAEETRKLMPVCVETKAIVS  TIQRKYKGIKIQEGVVDYGARFYFYTSKTTVASLINTLNDLNETLVTMPLGYVTHGLNLEEAARYMRSLK  VPATVSVSSPDAVTAYNGYLTSSSKTPEEHFIETISLAGSYKDWSYSGQSTQLGIEFLKRGDKSVYYTSN  PTTFHLDGEVITFDNLKTLLSLREVRTIKVFTTVDNINLHTQVVDMSMTYGQQFGPTYLDGADVTKIKPH  NSHEGKTFYVLPNDDTLRVEAFEYYHTTDPSFLGRYMSALNHTKKWKYPQVNGLTSIKWADNNCYLATAL  LTLQQIELKFNPPALQDAYYRARAGEAANFCALILAYCNKTVGELGDVRETMSYLFQHANLDSCKRVLNV  VCKTCGQQQTTLKGVEAVMYMGTLSYEQFKKGVQIPCTCGKQATKYLVQQESPFVMMSAPPAQYELKHGT  FTCASEYTGNYQCGHYKHITSKETLYCIDGALLTKSSEYKGPITDVFYKENSYTTTIKPVTYKLDGVVCT  EIDPKLDNYYKKDNSYFTEQPIDLVPNQPYPNASFDNFKFVCDNIKFADDLNQLTGYKKPASRELKVTFF  PDLNGDVVAIDYKHYTPSFKKGAKLLHKPIVWHVNNATNKATYKPNTWCIRCLWSTKPVETSNSFDVLKS  EDAQGMDNLACEDLKPVSEEVVENPTIQKDVLECNVKTTEVVGDIILKPANNSLKITEEVGHTDLMAAYV  DNSSLTIKKPNELSRVLGLKTLATHGLAAVNSVPWDTIANYAKPFLNKVVSTTTNIVTRCLNRVCTNYMP  YFFTLLLQLCTFTRSTNSRIKASMPTTIAKNTVKSVGKFCLEASFNYLKSPNFSKLINIIIWFLLLSVCL  GSLIYSTAALGVLMSNLGMPSYCTGYREGYLNSTNVTIATYCTGSIPCSVCLSGLDSLDTYPSLETIQIT  ISSFKWDLTAFGLVAEWFLAYILFTRFFYVLGLAAIMQLFFSYFAVHFISNSWLMWLIINLVQMAPISAM  VRMYIFFASFYYVWKSYVHVVDGCNSSTCMMCYKRNRATRVECTTIVNGVRRSFYVYANGGKGFCKLHNW  NCVNCDTFCAGSTFISDEVARDLSLQFKRPINPTDQSSYIVDSVTVKNGSIHLYFDKAGQKTYERHSLSH  FVNLDNLRANNTKGSLPINVIVFDGKSKCEESSAKSASVYYSQLMCQPILLLDQALVSDVGDSAEVAVKM  FDAYVNTFSSTFNVPMEKLKTLVATAEAELAKNVSLDNVLSTFISAARQGFVDSDVETKDVVECLKLSHQ  SDIEVTGDSCNNYMLTYNKVENMTPRDLGACIDCSARHINAQVAKSHNIALIWNVKDFMSLSEQLRKQIR  SAAKKNNLPFKLTCATTRQVVNVVTTKIALKGGKIVNNWLKQLIKVTLVFLFVAAIFYLITPVHVMSKHT  DFSSEIIGYKAIDGGVTRDIASTDTCFANKHADFDTWFSQRGGSYTNDKACPLIAAVITREVGFVVPGLP  GTILRTTNGDFLHFLPRVFSAVGNICYTPSKLIEYTDFATSACVLAAECTIFKDASGKPVPYCYDTNVLE  GSVAYESLRPDTRYVLMDGSIIQFPNTYLEGSVRVVTTFDSEYCRHGTCERSEAGVCVSTSGRWVLNNDY  YRSLPGVFCGVDAVNLLTNMFTPLIQPIGALDISASIVAGGIVAIVVTCLAYYFMRFRRAFGEYSHVVAF  NTLLFLMSFTVLCLTPVYSFLPGVYSVIYLYLTFYLTNDVSFLAHIQWMVMFTPLVPFWITIAYIICIST  KHFYWFFSNYLKRRVVFNGVSFSTFEEAALCTFLLNKEMYLKLRSDVLLPLTQYNRYLALYNKYKYFSGA  MDTTSYREAACCHLAKALNDFSNSGSDVLYQPPQTSITSAVLQSGFRKMAFPSGKVEGCMVQVTCGTTTL  NGLWLDDVVYCPRHVICTSEDMLNPNYEDLLIRKSNHNFLVQAGNVQLRVIGHSMQNCVLKLKVDTANPK  TPKYKFVRIQPGQTFSVLACYNGSPSGVYQCAMRPNFTIKGSFLNGSCGSVGFNIDYDCVSFCYMHHMEL  PTGVHAGTDLEGNFYGPFVDRQTAQAAGTDTTITVNVLAWLYAAVINGDRWFLNRFTTTLNDFNLVAMKY  NYEPLTQDHVDILGPLSAQTGIAVLDMCASLKELLQNGMNGRTILGSALLEDEFTPFDVVRQCSGVTFQS  AVKRTIKGTHHWLLLTILTSLLVLVQSTQWSLFFFLYENAFLPFAMGIIAMSAFAMMFVKHKHAFLCLFL  LPSLATVAYFNMVYMPASWVMRIMTWLDMVDTSLSGFKLKDCVMYASAVVLLILMTARTVYDDGARRVWT  LMNVLTLVYKVYYGNALDQAISMWALIISVTSNYSGVVTTVMFLARGIVFMCVEYCPIFFITGNTLQCIM  LVYCFLGYFCTCYFGLFCLLNRYFRLTLGVYDYLVSTQEFRYMNSQGLLPPKNSIDAFKLNIKLLGVGGK  PCIKVATVQSKMSDVKCTSVVLLSVLQQLRVESSSKLWAQCVQLHNDILLAKDTTEAFEKMVSLLSVLLS  MQGAVDINKLCEEMLDNRATLQAIASEFSSLPSYAAFATAQEAYEQAVANGDSEVVLKKLKKSLNVAKSE  FDRDAAMQRKLEKMADQAMTQMYKQARSEDKRAKVTSAMQTMLFTMLRKLDNDALNNIINNARDGCVPLN  IIPLTTAAKLMVVIPDYNTYKNTCDGTTFTYASALWEIQQVVDADSKIVQLSEISMDNSPNLAWPLIVTA  LRANSAVKLQNNELSPVALRQMSCAAGTTQTACTDDNALAYYNTTKGGRFVLALLSDLQDLKWARFPKSD  GTGTIYTELEPPCRFVTDTPKGPKVKYLYFIKGLNNLNRGMVLGSLAATVRLQAGNATEVPANSTVLSFC  AFAVDAAKAYKDYLASGGQPITNCVKMLCTHTGTGQAITVTPEANMDQESFGGASCCLYCRCHIDHPNPK  GFCDLKGKYVQIPTTCANDPVGFTLKNTVCTVCGMWKGYGCSCDQLREPMLQSADAQSFLNGFAV |
| **sp\|P0DTD1\|1ab** | Replicase polyprotein 1ab | >sp\|P0DTD1.1\|R1AB_SARS2  MESLVPGFNEKTHVQLSLPVLQVRDVLVRGFGDSVEEVLSEARQHLKDGTCGLVEVEKGVLPQLEQPYVF  IKRSDARTAPHGHVMVELVAELEGIQYGRSGETLGVLVPHVGEIPVAYRKVLLRKNGNKGAGGHSYGADL  KSFDLGDELGTDPYEDFQENWNTKHSSGVTRELMRELNGGAYTRYVDNNFCGPDGYPLECIKDLLARAGK  ASCTLSEQLDFIDTKRGVYCCREHEHEIAWYTERSEKSYELQTPFEIKLAKKFDTFNGECPNFVFPLNSI  IKTIQPRVEKKKLDGFMGRIRSVYPVASPNECNQMCLSTLMKCDHCGETSWQTGDFVKATCEFCGTENLT  KEGATTCGYLPQNAVVKIYCPACHNSEVGPEHSLAEYHNESGLKTILRKGGRTIAFGGCVFSYVGCHNKC  AYWVPRASANIGCNHTGVVGEGSEGLNDNLLEILQKEKVNINIVGDFKLNEEIAIILASFSASTSAFVET  VKGLDYKAFKQIVESCGNFKVTKGKAKKGAWNIGEQKSILSPLYAFASEAARVVRSIFSRTLETAQNSVR  VLQKAAITILDGISQYSLRLIDAMMFTSDLATNNLVVMAYITGGVVQLTSQWLTNIFGTVYEKLKPVLDW  LEEKFKEGVEFLRDGWEIVKFISTCACEIVGGQIVTCAKEIKESVQTFFKLVNKFLALCADSIIIGGAKL  KALNLGETFVTHSKGLYRKCVKSREETGLLMPLKAPKEIIFLEGETLPTEVLTEEVVLKTGDLQPLEQPT  SEAVEAPLVGTPVCINGLMLLEIKDTEKYCALAPNMMVTNNTFTLKGGAPTKVTFGDDTVIEVQGYKSVN  ITFELDERIDKVLNEKCSAYTVELGTEVNEFACVVADAVIKTLQPVSELLTPLGIDLDEWSMATYYLFDE  SGEFKLASHMYCSFYPPDEDEEEGDCEEEEFEPSTQYEYGTEDDYQGKPLEFGATSAALQPEEEQEEDWL  DDDSQQTVGQQDGSEDNQTTTIQTIVEVQPQLEMELTPVVQTIEVNSFSGYLKLTDNVYIKNADIVEEAK  KVKPTVVVNAANVYLKHGGGVAGALNKATNNAMQVESDDYIATNGPLKVGGSCVLSGHNLAKHCLHVVGP  NVNKGEDIQLLKSAYENFNQHEVLLAPLLSAGIFGADPIHSLRVCVDTVRTNVYLAVFDKNLYDKLVSSF  LEMKSEKQVEQKIAEIPKEEVKPFITESKPSVEQRKQDDKKIKACVEEVTTTLEETKFLTENLLLYIDIN  GNLHPDSATLVSDIDITFLKKDAPYIVGDVVQEGVLTAVVIPTKKAGGTTEMLAKALRKVPTDNYITTYP  GQGLNGYTVEEAKTVLKKCKSAFYILPSIISNEKQEILGTVSWNLREMLAHAEETRKLMPVCVETKAIVS  TIQRKYKGIKIQEGVVDYGARFYFYTSKTTVASLINTLNDLNETLVTMPLGYVTHGLNLEEAARYMRSLK  VPATVSVSSPDAVTAYNGYLTSSSKTPEEHFIETISLAGSYKDWSYSGQSTQLGIEFLKRGDKSVYYTSN  PTTFHLDGEVITFDNLKTLLSLREVRTIKVFTTVDNINLHTQVVDMSMTYGQQFGPTYLDGADVTKIKPH  NSHEGKTFYVLPNDDTLRVEAFEYYHTTDPSFLGRYMSALNHTKKWKYPQVNGLTSIKWADNNCYLATAL  LTLQQIELKFNPPALQDAYYRARAGEAANFCALILAYCNKTVGELGDVRETMSYLFQHANLDSCKRVLNV  VCKTCGQQQTTLKGVEAVMYMGTLSYEQFKKGVQIPCTCGKQATKYLVQQESPFVMMSAPPAQYELKHGT  FTCASEYTGNYQCGHYKHITSKETLYCIDGALLTKSSEYKGPITDVFYKENSYTTTIKPVTYKLDGVVCT  EIDPKLDNYYKKDNSYFTEQPIDLVPNQPYPNASFDNFKFVCDNIKFADDLNQLTGYKKPASRELKVTFF  PDLNGDVVAIDYKHYTPSFKKGAKLLHKPIVWHVNNATNKATYKPNTWCIRCLWSTKPVETSNSFDVLKS  EDAQGMDNLACEDLKPVSEEVVENPTIQKDVLECNVKTTEVVGDIILKPANNSLKITEEVGHTDLMAAYV  DNSSLTIKKPNELSRVLGLKTLATHGLAAVNSVPWDTIANYAKPFLNKVVSTTTNIVTRCLNRVCTNYMP  YFFTLLLQLCTFTRSTNSRIKASMPTTIAKNTVKSVGKFCLEASFNYLKSPNFSKLINIIIWFLLLSVCL  GSLIYSTAALGVLMSNLGMPSYCTGYREGYLNSTNVTIATYCTGSIPCSVCLSGLDSLDTYPSLETIQIT  ISSFKWDLTAFGLVAEWFLAYILFTRFFYVLGLAAIMQLFFSYFAVHFISNSWLMWLIINLVQMAPISAM  VRMYIFFASFYYVWKSYVHVVDGCNSSTCMMCYKRNRATRVECTTIVNGVRRSFYVYANGGKGFCKLHNW  NCVNCDTFCAGSTFISDEVARDLSLQFKRPINPTDQSSYIVDSVTVKNGSIHLYFDKAGQKTYERHSLSH  FVNLDNLRANNTKGSLPINVIVFDGKSKCEESSAKSASVYYSQLMCQPILLLDQALVSDVGDSAEVAVKM  FDAYVNTFSSTFNVPMEKLKTLVATAEAELAKNVSLDNVLSTFISAARQGFVDSDVETKDVVECLKLSHQ  SDIEVTGDSCNNYMLTYNKVENMTPRDLGACIDCSARHINAQVAKSHNIALIWNVKDFMSLSEQLRKQIR  SAAKKNNLPFKLTCATTRQVVNVVTTKIALKGGKIVNNWLKQLIKVTLVFLFVAAIFYLITPVHVMSKHT  DFSSEIIGYKAIDGGVTRDIASTDTCFANKHADFDTWFSQRGGSYTNDKACPLIAAVITREVGFVVPGLP  GTILRTTNGDFLHFLPRVFSAVGNICYTPSKLIEYTDFATSACVLAAECTIFKDASGKPVPYCYDTNVLE  GSVAYESLRPDTRYVLMDGSIIQFPNTYLEGSVRVVTTFDSEYCRHGTCERSEAGVCVSTSGRWVLNNDY  YRSLPGVFCGVDAVNLLTNMFTPLIQPIGALDISASIVAGGIVAIVVTCLAYYFMRFRRAFGEYSHVVAF  NTLLFLMSFTVLCLTPVYSFLPGVYSVIYLYLTFYLTNDVSFLAHIQWMVMFTPLVPFWITIAYIICIST  KHFYWFFSNYLKRRVVFNGVSFSTFEEAALCTFLLNKEMYLKLRSDVLLPLTQYNRYLALYNKYKYFSGA  MDTTSYREAACCHLAKALNDFSNSGSDVLYQPPQTSITSAVLQSGFRKMAFPSGKVEGCMVQVTCGTTTL  NGLWLDDVVYCPRHVICTSEDMLNPNYEDLLIRKSNHNFLVQAGNVQLRVIGHSMQNCVLKLKVDTANPK  TPKYKFVRIQPGQTFSVLACYNGSPSGVYQCAMRPNFTIKGSFLNGSCGSVGFNIDYDCVSFCYMHHMEL  PTGVHAGTDLEGNFYGPFVDRQTAQAAGTDTTITVNVLAWLYAAVINGDRWFLNRFTTTLNDFNLVAMKY  NYEPLTQDHVDILGPLSAQTGIAVLDMCASLKELLQNGMNGRTILGSALLEDEFTPFDVVRQCSGVTFQS  AVKRTIKGTHHWLLLTILTSLLVLVQSTQWSLFFFLYENAFLPFAMGIIAMSAFAMMFVKHKHAFLCLFL  LPSLATVAYFNMVYMPASWVMRIMTWLDMVDTSLSGFKLKDCVMYASAVVLLILMTARTVYDDGARRVWT  LMNVLTLVYKVYYGNALDQAISMWALIISVTSNYSGVVTTVMFLARGIVFMCVEYCPIFFITGNTLQCIM  LVYCFLGYFCTCYFGLFCLLNRYFRLTLGVYDYLVSTQEFRYMNSQGLLPPKNSIDAFKLNIKLLGVGGK  PCIKVATVQSKMSDVKCTSVVLLSVLQQLRVESSSKLWAQCVQLHNDILLAKDTTEAFEKMVSLLSVLLS  MQGAVDINKLCEEMLDNRATLQAIASEFSSLPSYAAFATAQEAYEQAVANGDSEVVLKKLKKSLNVAKSE  FDRDAAMQRKLEKMADQAMTQMYKQARSEDKRAKVTSAMQTMLFTMLRKLDNDALNNIINNARDGCVPLN  IIPLTTAAKLMVVIPDYNTYKNTCDGTTFTYASALWEIQQVVDADSKIVQLSEISMDNSPNLAWPLIVTA  LRANSAVKLQNNELSPVALRQMSCAAGTTQTACTDDNALAYYNTTKGGRFVLALLSDLQDLKWARFPKSD  GTGTIYTELEPPCRFVTDTPKGPKVKYLYFIKGLNNLNRGMVLGSLAATVRLQAGNATEVPANSTVLSFC  AFAVDAAKAYKDYLASGGQPITNCVKMLCTHTGTGQAITVTPEANMDQESFGGASCCLYCRCHIDHPNPK  GFCDLKGKYVQIPTTCANDPVGFTLKNTVCTVCGMWKGYGCSCDQLREPMLQSADAQSFLNRVCGVSAAR  LTPCGTGTSTDVVYRAFDIYNDKVAGFAKFLKTNCCRFQEKDEDDNLIDSYFVVKRHTFSNYQHEETIYN  LLKDCPAVAKHDFFKFRIDGDMVPHISRQRLTKYTMADLVYALRHFDEGNCDTLKEILVTYNCCDDDYFN  KKDWYDFVENPDILRVYANLGERVRQALLKTVQFCDAMRNAGIVGVLTLDNQDLNGNWYDFGDFIQTTPG  SGVPVVDSYYSLLMPILTLTRALTAESHVDTDLTKPYIKWDLLKYDFTEERLKLFDRYFKYWDQTYHPNC  VNCLDDRCILHCANFNVLFSTVFPPTSFGPLVRKIFVDGVPFVVSTGYHFRELGVVHNQDVNLHSSRLSF  KELLVYAADPAMHAASGNLLLDKRTTCFSVAALTNNVAFQTVKPGNFNKDFYDFAVSKGFFKEGSSVELK  HFFFAQDGNAAISDYDYYRYNLPTMCDIRQLLFVVEVVDKYFDCYDGGCINANQVIVNNLDKSAGFPFNK  WGKARLYYDSMSYEDQDALFAYTKRNVIPTITQMNLKYAISAKNRARTVAGVSICSTMTNRQFHQKLLKS  IAATRGATVVIGTSKFYGGWHNMLKTVYSDVENPHLMGWDYPKCDRAMPNMLRIMASLVLARKHTTCCSL  SHRFYRLANECAQVLSEMVMCGGSLYVKPGGTSSGDATTAYANSVFNICQAVTANVNALLSTDGNKIADK  YVRNLQHRLYECLYRNRDVDTDFVNEFYAYLRKHFSMMILSDDAVVCFNSTYASQGLVASIKNFKSVLYY  QNNVFMSEAKCWTETDLTKGPHEFCSQHTMLVKQGDDYVYLPYPDPSRILGAGCFVDDIVKTDGTLMIER  FVSLAIDAYPLTKHPNQEYADVFHLYLQYIRKLHDELTGHMLDMYSVMLTNDNTSRYWEPEFYEAMYTPH  TVLQAVGACVLCNSQTSLRCGACIRRPFLCCKCCYDHVISTSHKLVLSVNPYVCNAPGCDVTDVTQLYLG  GMSYYCKSHKPPISFPLCANGQVFGLYKNTCVGSDNVTDFNAIATCDWTNAGDYILANTCTERLKLFAAE  TLKATEETFKLSYGIATVREVLSDRELHLSWEVGKPRPPLNRNYVFTGYRVTKNSKVQIGEYTFEKGDYG  DAVVYRGTTTYKLNVGDYFVLTSHTVMPLSAPTLVPQEHYVRITGLYPTLNISDEFSSNVANYQKVGMQK  YSTLQGPPGTGKSHFAIGLALYYPSARIVYTACSHAAVDALCEKALKYLPIDKCSRIIPARARVECFDKF  KVNSTLEQYVFCTVNALPETTADIVVFDEISMATNYDLSVVNARLRAKHYVYIGDPAQLPAPRTLLTKGT  LEPEYFNSVCRLMKTIGPDMFLGTCRRCPAEIVDTVSALVYDNKLKAHKDKSAQCFKMFYKGVITHDVSS  AINRPQIGVVREFLTRNPAWRKAVFISPYNSQNAVASKILGLPTQTVDSSQGSEYDYVIFTQTTETAHSC  NVNRFNVAITRAKVGILCIMSDRDLYDKLQFTSLEIPRRNVATLQAENVTGLFKDCSKVITGLHPTQAPT  HLSVDTKFKTEGLCVDIPGIPKDMTYRRLISMMGFKMNYQVNGYPNMFITREEAIRHVRAWIGFDVEGCH  ATREAVGTNLPLQLGFSTGVNLVAVPTGYVDTPNNTDFSRVSAKPPPGDQFKHLIPLMYKGLPWNVVRIK  IVQMLSDTLKNLSDRVVFVLWAHGFELTSMKYFVKIGPERTCCLCDRRATCFSTASDTYACWHHSIGFDY  VYNPFMIDVQQWGFTGNLQSNHDLYCQVHGNAHVASCDAIMTRCLAVHECFVKRVDWTIEYPIIGDELKI  NAACRKVQHMVVKAALLADKFPVLHDIGNPKAIKCVPQADVEWKFYDAQPCSDKAYKIEELFYSYATHSD  KFTDGVCLFWNCNVDRYPANSIVCRFDTRVLSNLNLPGCDGGSLYVNKHAFHTPAFDKSAFVNLKQLPFF  YYSDSPCESHGKQVVSDIDYVPLKSATCITRCNLGGAVCRHHANEYRLYLDAYNMMISAGFSLWVYKQFD  TYNLWNTFTRLQSLENVAFNVVNKGHFDGQQGEVPVSIINNTVYTKVDGVDVELFENKTTLPVNVAFELW  AKRNIKPVPEVKILNNLGVDIAANTVIWDYKRDAPAHISTIGVCSMTDIAKKPTETICAPLTVFFDGRVD  GQVDLFRNARNGVLITEGSVKGLQPSVGPKQASLNGVTLIGEAVKTQFNYYKKVDGVVQQLPETYFTQSR  NLQEFKPRSQMEIDFLELAMDEFIERYKLEGYAFEHIVYGDFSHSQLGGLHLLIGLAKRFKESPFELEDF  IPMDSTVKNYFITDAQTGSSKCVCSVIDLLLDDFVEIIKSQDLSVVSKVVKVTIDYTEISFMLWCKDGHV  ETFYPKLQSSQAWQPGVAMPNLYKMQRMLLEKCDLQNYGDSATLPKGIMMNVAKYTQLCQYLNTLTLAVP  YNMRVIHFGAGSDKGVAPGTAVLRQWLPTGTLLVDSDLNDFVSDADSTLIGDCATVHTANKWDLIISDMY  DPKTKNVTKENDSKEGFFTYICGFIQQKLALGGSVAIKITEHSWNADLYKLMGHFAWWTAFVTNVNASSS  EAFLIGCNYLGKPREQIDGYVMHANYIFWRNTNPIQLSSYSLFDMSKFPLKLRGTAVMSLKEGQINDMIL  SLLSKGRLIIRENNRVVISSDVLVNN |
| **YP_009742609.1** | Nsp2 | >YP_009742609.1 nsp2 [Severe acute respiratory syndrome coronavirus 2]  AYTRYVDNNFCGPDGYPLECIKDLLARAGKASCTLSEQLDFIDTKRGVYCCREHEHEIAWYTERSEKSYE  LQTPFEIKLAKKFDTFNGECPNFVFPLNSIIKTIQPRVEKKKLDGFMGRIRSVYPVASPNECNQMCLSTL  MKCDHCGETSWQTGDFVKATCEFCGTENLTKEGATTCGYLPQNAVVKIYCPACHNSEVGPEHSLAEYHNE  SGLKTILRKGGRTIAFGGCVFSYVGCHNKCAYWVPRASANIGCNHTGVVGEGSEGLNDNLLEILQKEKVN  INIVGDFKLNEEIAIILASFSASTSAFVETVKGLDYKAFKQIVESCGNFKVTKGKAKKGAWNIGEQKSIL  SPLYAFASEAARVVRSIFSRTLETAQNSVRVLQKAAITILDGISQYSLRLIDAMMFTSDLATNNLVVMAY  ITGGVVQLTSQWLTNIFGTVYEKLKPVLDWLEEKFKEGVEFLRDGWEIVKFISTCACEIVGGQIVTCAKE  IKESVQTFFKLVNKFLALCADSIIIGGAKLKALNLGETFVTHSKGLYRKCVKSREETGLLMPLKAPKEII  FLEGETLPTEVLTEEVVLKTGDLQPLEQPTSEAVEAPLVGTPVCINGLMLLEIKDTEKYCALAPNMMVTN  NTFTLKGG |
| **YP_009742611.1** | Nsp4 | >YP_009742611.1 nsp4 [Severe acute respiratory syndrome coronavirus 2]  KIVNNWLKQLIKVTLVFLFVAAIFYLITPVHVMSKHTDFSSEIIGYKAIDGGVTRDIASTDTCFANKHAD  FDTWFSQRGGSYTNDKACPLIAAVITREVGFVVPGLPGTILRTTNGDFLHFLPRVFSAVGNICYTPSKLI  EYTDFATSACVLAAECTIFKDASGKPVPYCYDTNVLEGSVAYESLRPDTRYVLMDGSIIQFPNTYLEGSV  RVVTTFDSEYCRHGTCERSEAGVCVSTSGRWVLNNDYYRSLPGVFCGVDAVNLLTNMFTPLIQPIGALDI  SASIVAGGIVAIVVTCLAYYFMRFRRAFGEYSHVVAFNTLLFLMSFTVLCLTPVYSFLPGVYSVIYLYLT  FYLTNDVSFLAHIQWMVMFTPLVPFWITIAYIICISTKHFYWFFSNYLKRRVVFNGVSFSTFEEAALCTF  LLNKEMYLKLRSDVLLPLTQYNRYLALYNKYKYFSGAMDTTSYREAACCHLAKALNDFSNSGSDVLYQPP  QTSITSAVLQ |
| **YP_009742610.1** | Nsp 3 | >YP_009742610.1 nsp3 [Severe acute respiratory syndrome coronavirus 2]  APTKVTFGDDTVIEVQGYKSVNITFELDERIDKVLNEKCSAYTVELGTEVNEFACVVADAVIKTLQPVSE  LLTPLGIDLDEWSMATYYLFDESGEFKLASHMYCSFYPPDEDEEEGDCEEEEFEPSTQYEYGTEDDYQGK  PLEFGATSAALQPEEEQEEDWLDDDSQQTVGQQDGSEDNQTTTIQTIVEVQPQLEMELTPVVQTIEVNSF  SGYLKLTDNVYIKNADIVEEAKKVKPTVVVNAANVYLKHGGGVAGALNKATNNAMQVESDDYIATNGPLK  VGGSCVLSGHNLAKHCLHVVGPNVNKGEDIQLLKSAYENFNQHEVLLAPLLSAGIFGADPIHSLRVCVDT  VRTNVYLAVFDKNLYDKLVSSFLEMKSEKQVEQKIAEIPKEEVKPFITESKPSVEQRKQDDKKIKACVEE  VTTTLEETKFLTENLLLYIDINGNLHPDSATLVSDIDITFLKKDAPYIVGDVVQEGVLTAVVIPTKKAGG  TTEMLAKALRKVPTDNYITTYPGQGLNGYTVEEAKTVLKKCKSAFYILPSIISNEKQEILGTVSWNLREM  LAHAEETRKLMPVCVETKAIVSTIQRKYKGIKIQEGVVDYGARFYFYTSKTTVASLINTLNDLNETLVTM  PLGYVTHGLNLEEAARYMRSLKVPATVSVSSPDAVTAYNGYLTSSSKTPEEHFIETISLAGSYKDWSYSG  QSTQLGIEFLKRGDKSVYYTSNPTTFHLDGEVITFDNLKTLLSLREVRTIKVFTTVDNINLHTQVVDMSM  TYGQQFGPTYLDGADVTKIKPHNSHEGKTFYVLPNDDTLRVEAFEYYHTTDPSFLGRYMSALNHTKKWKY  PQVNGLTSIKWADNNCYLATALLTLQQIELKFNPPALQDAYYRARAGEAANFCALILAYCNKTVGELGDV  RETMSYLFQHANLDSCKRVLNVVCKTCGQQQTTLKGVEAVMYMGTLSYEQFKKGVQIPCTCGKQATKYLV  QQESPFVMMSAPPAQYELKHGTFTCASEYTGNYQCGHYKHITSKETLYCIDGALLTKSSEYKGPITDVFY  KENSYTTTIKPVTYKLDGVVCTEIDPKLDNYYKKDNSYFTEQPIDLVPNQPYPNASFDNFKFVCDNIKFA  DDLNQLTGYKKPASRELKVTFFPDLNGDVVAIDYKHYTPSFKKGAKLLHKPIVWHVNNATNKATYKPNTW  CIRCLWSTKPVETSNSFDVLKSEDAQGMDNLACEDLKPVSEEVVENPTIQKDVLECNVKTTEVVGDIILK  PANNSLKITEEVGHTDLMAAYVDNSSLTIKKPNELSRVLGLKTLATHGLAAVNSVPWDTIANYAKPFLNK  VVSTTTNIVTRCLNRVCTNYMPYFFTLLLQLCTFTRSTNSRIKASMPTTIAKNTVKSVGKFCLEASFNYL  KSPNFSKLINIIIWFLLLSVCLGSLIYSTAALGVLMSNLGMPSYCTGYREGYLNSTNVTIATYCTGSIPC  SVCLSGLDSLDTYPSLETIQITISSFKWDLTAFGLVAEWFLAYILFTRFFYVLGLAAIMQLFFSYFAVHF  ISNSWLMWLIINLVQMAPISAMVRMYIFFASFYYVWKSYVHVVDGCNSSTCMMCYKRNRATRVECTTIVN  GVRRSFYVYANGGKGFCKLHNWNCVNCDTFCAGSTFISDEVARDLSLQFKRPINPTDQSSYIVDSVTVKN  GSIHLYFDKAGQKTYERHSLSHFVNLDNLRANNTKGSLPINVIVFDGKSKCEESSAKSASVYYSQLMCQP  ILLLDQALVSDVGDSAEVAVKMFDAYVNTFSSTFNVPMEKLKTLVATAEAELAKNVSLDNVLSTFISAAR  QGFVDSDVETKDVVECLKLSHQSDIEVTGDSCNNYMLTYNKVENMTPRDLGACIDCSARHINAQVAKSHN  IALIWNVKDFMSLSEQLRKQIRSAAKKNNLPFKLTCATTRQVVNVVTTKIALKGG |
| **sp\|P0DTC6\|** | nsp 6 | >sp\|P0DTC6.1\|NS6_SARS2 RecName: Full=ORF6 protein; Short=ORF6; AltName: Full=Accessory protein 6; AltName: Full=Non-structural protein 6; Short=ns6; AltName: Full=Protein X3  MFHLVDFQVTIAEILLIIMRTFKVSIWNLDYIINLIIKNLSKSLTENKYSQLDEEQPMEID |
| **YP_009742614.1** | Nsp 7 | >YP_009742614.1 nsp7 [Severe acute respiratory syndrome coronavirus 2]  SKMSDVKCTSVVLLSVLQQLRVESSSKLWAQCVQLHNDILLAKDTTEAFEKMVSLLSVLLSMQGAVDINK  LCEEMLDNRATLQ |
| **sp\|P0DTC8\|** | nsp 8 | >sp\|P0DTC8.1\|NS8_SARS2 RecName: Full=ORF8 protein; Short=ORF8; AltName: Full=Non-structural protein 8; Short=ns8; Flags: Precursor  MKFLVFLGIITTVAAFHQECSLQSCTQHQPYVVDDPCPIHFYSKWYIRVGARKSAPLIELCVDEAGSKSP  IQYIDIGNYTVSCLPFTINCQEPKLGSLVVRCSFYEDFLEYHDVRVVLDFI |
| **YP_009725312.1** | nsp11 | >YP_009725312.1 nsp11 [Severe acute respiratory syndrome coronavirus 2]  SADAQSFLNGFAV |
| **YP_009742616.1** | Nsp 9 | >YP_009742616.1 nsp9 [Severe acute respiratory syndrome coronavirus 2]  NNELSPVALRQMSCAAGTTQTACTDDNALAYYNTTKGGRFVLALLSDLQDLKWARFPKSDGTGTIYTELE  PPCRFVTDTPKGPKVKYLYFIKGLNNLNRGMVLGSLAATVRLQ |
| **YP_009742617.1** | Nsp 10 | >YP_009742617.1 nsp10 [Severe acute respiratory syndrome coronavirus 2]  AGNATEVPANSTVLSFCAFAVDAAKAYKDYLASGGQPITNCVKMLCTHTGTGQAITVTPEANMDQESFGG  ASCCLYCRCHIDHPNPKGFCDLKGKYVQIPTTCANDPVGFTLKNTVCTVCGMWKGYGCSCDQLREPMLQ |
| **6M71_A** | NSP 12 | SADAQSFLNRVCGVSAARLTPCGTGTSTDVVYRAFDIYNDKVAGFAKFLKTNCCRFQEKDEDDNLIDSYFVVKRHTFSNYQHEETIYNLLKDCPAVAKHDFFKFRIDGDMVPHISRQRLTKYTMADLVYALRHFDEGNCDTLKEILVTYNCCDDDYFNKKDWYDFVENPDILRVYANLGERVRQALLKTVQFCDAMRNAGIVGVLTLDNQDLNGNWYDFGDFIQTTPGSGVPVVDSYYSLLMPILTLTRALTAESHVDTDLTKPYIKWDLLKYDFTEERLKLFDRYFKYWDQTYHPNCVNCLDDRCILHCANFNVLFSTVFPPTSFGPLVRKIFVDGVPFVVSTGYHFRELGVVHNQDVNLHSSRLSFKELLVYAADPAMHAASGNLLLDKRTTCFSVAALTNNVAFQTVKPGNFNKDFYDFAVSKGFFKEGSSVELKHFFFAQDGNAAISDYDYYRYNLPTMCDIRQLLFVVEVVDKYFDCYDGGCINANQVIVNNLDKSAGFPFNKWGKARLYYDSMSYEDQDALFAYTKRNVIPTITQMNLKYAISAKNRARTVAGVSICSTMTNRQFHQKLLKSIAATRGATVVIGTSKFYGGWHNMLKTVYSDVENPHLMGWDYPKCDRAMPNMLRIMASLVLARKHTTCCSLSHRFYRLANECAQVLSEMVMCGGSLYVKPGGTSSGDATTAYANSVFNICQAVTANVNALLSTDGNKIADKYVRNLQHRLYECLYRNRDVDTDFVNEFYAYLRKHFSMMILSDDAVVCFNSTYASQGLVASIKNFKSVLYYQNNVFMSEAKCWTETDLTKGPHEFCSQHTMLVKQGDDYVYLPYPDPSRILGAGCFVDDIVKTDGTLMIERFVSLAIDAYPLTKHPNQEYADVFHLYLQYIRKLHDELTGHMLDMYSVMLTNDNTSRYWEPEFYEAMYTPHTVLQHHHHHHHHHH |
