## Supplementary material for "Inferring Toll-Like Receptor induced epitope subunit vaccine candidate against SARS-CoV-2: A Reverse Vaccinology approach": Table s2

**TABLE S3: HTL and CTL epitope with their respective antigenic, allergenic and conservancy score.**

| **peptide** | **protein** | **Similarity index** | **Antigenicity** | **Alleargenicity** | **Conservancy** |
| --- | --- | --- | --- | --- | --- |
| AIILASFSASTSAFV | **NSP 2** | 0.65 | Probable antigen | NON-ALLERGEN | 100% |
| ERSEKSYELQ | **NSP 2** | 0.67 | Probable antigen | NON-ALLERGEN | 100% |
| ESVQTFFKL | **NSP 2** | 0.72 | Probable antigen | NON-ALLERGEN | 100% |
| FMGRIRSVY | **NSP 2** | 0.72 | Probable antigen | NON-ALLERGEN | 100% |
| GVVGEGSEGL | **NSP 2** | 0.68 | Probable antigen | NON-ALLERGEN | 100% |
| HNSEVGPEHSLAEYHNES | **NSP 2** | 0.65 | Probable antigen | NON-ALLERGEN | 100% |
| ILASFSASTSAFVET | **NSP 2** | 0.68 | Probable antigen | NON-ALLERGEN | 100% |
| KLAKKFDTF | **NSP 2** | 0.73 | Probable antigen | NON-ALLERGEN | 100% |
| LQPLEQPTSEAVEAPLV | **NSP 2** | 0.57 | Probable antigen | NON-ALLERGEN | 100% |
| LSPLYAFASEAARVV | **NSP 2** | 0.58 | Probable antigen | NON-ALLERGEN | 100% |
| LVNKFLALCADSIII | **NSP 2** | 0.65 | Probable antigen | NON-ALLERGEN | 100% |
| LYAFASEAARVVRSI | **NSP 2** | 0.61 | Probable antigen | NON-ALLERGEN | 100% |
| NLTKEGATTCGYL | **NSP 2** | 0.62 | Probable antigen | NON-ALLERGEN | 100% |
| NMMVTNNTF | **NSP 2** | 0.76 | Probable antigen | NON-ALLERGEN | 100% |
| PLYAFASEAARVVRS | **NSP 2** | 0.57 | Probable antigen | NON-ALLERGEN | 100% |
| PRVEKK | **NSP 2** | 0.77 | Probable antigen | NON-ALLERGEN | 100% |
| RAGKASCTL | **NSP 2** | 0.67 | Probable antigen | NON-ALLERGEN | 100% |
| TEKYCALAPNMMVTN | **NSP 2** | 0.6 | Probable antigen | NON-ALLERGEN | 100% |
| TILDGISQY | **NSP 2** | 0.75 | Probable antigen | NON-ALLERGEN | 100% |
| TLPTE | **NSP 2** | 0.82 | Probable antigen | NON-ALLERGEN | 100% |
| TSQWLTNIF | **NSP 2** | 0.76 | Probable antigen | NON-ALLERGEN | 100% |
| VTKGKAKKGAWNIG | **NSP 2** | 0.6 | Probable antigen | NON-ALLERGEN | 100% |
| AAIFYLITPVHVMSK | **NSP 4** | 0.66 | Probable antigen | NON-ALLERGEN | 100% |
| AMDTTSYR | **NSP 4** | 0.75 | Probable antigen | NON-ALLERGEN | 100% |
| DFSNSGSDVLYQPPQTSI | **NSP 4** | 0.68 | Probable antigen | NON-ALLERGEN | 100% |
| ESLRPDTRY | **NSP 4** | 0.69 | Probable antigen | NON-ALLERGEN | 100% |
| FEEAALCTF | **NSP 4** | 0.71 | Probable antigen | NON-ALLERGEN | 100% |
| FMRFRRAFGEYSHVV | **NSP 4** | 0.58 | Probable antigen | NON-ALLERGEN | 100% |
| FYWFFSNYL | **NSP 4** | 0.76 | Probable antigen | NON-ALLERGEN | 100% |
| LMSFTVLCL | **NSP 4** | 0.68 | Probable antigen | NON-ALLERGEN | 100% |
| SNSGSDVLY | **NSP 4** | 0.73 | Probable antigen | NON-ALLERGEN | 100% |
| TRYVLMDGSIIQFPN | **NSP 4** | 0.6 | Probable antigen | NON-ALLERGEN | 100% |
| VIYLYLTFY | **NSP 4** | 0.73 | Probable antigen | NON-ALLERGEN | 100% |
| EAFEKMVSL | NSP 7 | 0.68 | Probable antigen | NON-ALLERGEN | 100% |
| LSVLLSMQGAVDINK | NSP 7 | 0.66 | Probable antigen | NON-ALLERGEN | 100% |
| TEAFEKMVSLLSVLL | NSP 7 | 0.6 | Probable antigen | NON-ALLERGEN | 100% |
| CSSGTYEGNSPFHPL | NSP 7a | 0.59 | Probable antigen | NON-ALLERGEN | 100% |
| VKHVYQLRARSVSPK | NSP 7a | 0.57 | Probable antigen | NON-ALLERGEN | 100% |
| AAGTTQTACTDDN | NSP 9 | 0.67 | Probable antigen | NON-ALLERGEN | 100% |
| FPKSDGTGTIYTELEP | NSP 9 | 0.56 | Probable antigen | NON-ALLERGEN | 100% |
| KYLYFIKGLNNLNRG | NSP 9 | 0.61 | Probable antigen | NON-ALLERGEN | 100% |
| RGMVLGSLAATVRLQ | NSP 9 | 0.61 | Probable antigen | NON-ALLERGEN | 100% |
| YLYFIKGLNNLNRGM | NSP 9 | 0.62 | Probable antigen | NON-ALLERGEN | 100% |
| DHPNPKGF | NSP 10 | 0.75 | Probable antigen | NON-ALLERGEN | 100% |
| FAVDAAKAY | NSP 10 | 0.74 | Probable antigen | NON-ALLERGEN | 100% |
| GNATEVPAN | NSP 10 | 0.71 | Probable antigen | NON-ALLERGEN | 100% |
| DDYVYLPYPDPSRI | NSP 12 | 0.61 | Probable antigen | NON-ALLERGEN | 100% |
| DSYYSLLMPILTLTR | NSP 12 | 0.63 | Probable antigen | NON-ALLERGEN | 100% |
| DVDTD | NSP 12 | 0.84 | Probable antigen | NON-ALLERGEN | 100% |
| ELLVYAADPAMHAAS | NSP 12 | 0.6 | Probable antigen | NON-ALLERGEN | 100% |
| ETDLTKGPHEF | NSP 12 | 0.64 | Probable antigen | NON-ALLERGEN | 100% |
| FVNEFYAYL | NSP 12 | 0.73 | Probable antigen | NON-ALLERGEN | 100% |
| FYAYLRKHFSMMILS | NSP 12 | 0.67 | Probable antigen | NON-ALLERGEN | 100% |
| HQKLLKSIAATRGAT | NSP 12 | 0.64 | Probable antigen | NON-ALLERGEN | 100% |
| IQTTPGSGVPVV | NSP 12 | 0.63 | Probable antigen | NON-ALLERGEN | 100% |
| KELLVYAADPAMHAA | NSP 12 | 0.59 | Probable antigen | NON-ALLERGEN | 100% |
| KLFDRYFKY | NSP 12 | 0.74 | Probable antigen | NON-ALLERGEN | 100% |
| KNFKSVLYY | NSP 12 | 0.75 | Probable antigen | NON-ALLERGEN | 100% |
| LKHFFFAQDGNAAIS | NSP 12 | 0.63 | Probable antigen | NON-ALLERGEN | 100% |
| LLDKRTTCF | NSP 12 | 0.7 | Probable antigen | NON-ALLERGEN | 100% |
| LLKSIAATRGATVVI | NSP 12 | 0.55 | Probable antigen | NON-ALLERGEN | 100% |
| LSFKELLVY | NSP 12 | 0.72 | Probable antigen | NON-ALLERGEN | 100% |
| LTKHPNQEY | NSP 12 | 0.7 | Probable antigen | NON-ALLERGEN | 100% |
| MLRIMASLVLARKHT | NSP 12 | 0.64 | Probable antigen | NON-ALLERGEN | 100% |
| MNLKYAISAKNRART | NSP 12 | 0.62 | Probable antigen | NON-ALLERGEN | 100% |
| PNMLRIMASLVLARK | NSP 12 | 0.58 | Probable antigen | NON-ALLERGEN | 100% |
| QKLLKSIAATRGATV | NSP 12 | 0.59 | Probable antigen | NON-ALLERGEN | 100% |
| QMNLKYAISAKNRAR | NSP 12 | 0.61 | Probable antigen | NON-ALLERGEN | 100% |
| SHRFYRLANECAQVL | NSP 12 | 0.59 | Probable antigen | NON-ALLERGEN | 100% |
| VKPGNFNK | NSP 12 | 0.72 | Probable antigen | NON-ALLERGEN | 100% |
| YFVVKRHTF | NSP 12 | 0.74 | Probable antigen | NON-ALLERGEN | 100% |
| AQKFNGLTVLPPLLT | SPIKE | 0.61 | Probable antigen | NON-ALLERGEN | 100% |
| ECSNLLLQY | SPIKE | 0.7 | Probable antigen | NON-ALLERGEN | 100% |
| FVSNGTHWF | SPIKE | 0.74 | Probable antigen | NON-ALLERGEN | 100% |
| GINITRFQTLLALHR | SPIKE | 0.64 | Probable antigen | NON-ALLERGEN | 100% |
| GWTFGAGAALQIPFA | SPIKE | 0.58 | Probable antigen | NON-ALLERGEN | 100% |
| IIAYTMSLGAENSVA | SPIKE | 0.64 | Probable antigen | NON-ALLERGEN | 100% |
| ILPDPSKPSKRS | SPIKE | 0.62 | Probable antigen | NON-ALLERGEN | 100% |
| ITRFQTLLALHRSYL | SPIKE | 0.65 | Probable antigen | NON-ALLERGEN | 100% |
| IYKTPPIKDF | SPIKE | 0.69 | Probable antigen | NON-ALLERGEN | 100% |
| KAHFP | SPIKE | 0.82 | Probable antigen | NON-ALLERGEN | 100% |
| KFNGLTVLPPLLTDE | SPIKE | 0.6 | Probable antigen | NON-ALLERGEN | 100% |
| KKFLPFQQF | SPIKE | 0.73 | Probable antigen | NON-ALLERGEN | 100% |
| KNHTSPDVDLG | SPIKE | 0.69 | Probable antigen | NON-ALLERGEN | 100% |
| LGQSKRVDF | SPIKE | 0.72 | Probable antigen | NON-ALLERGEN | 100% |
| LITGRLQSL | SPIKE | 0.71 | Probable antigen | NON-ALLERGEN | 100% |
| LTPGDSSSGWTAG | SPIKE | 0.61 | Probable antigen | NON-ALLERGEN | 100% |
| NATRFASVY | SPIKE | 0.74 | Probable antigen | NON-ALLERGEN | 100% |
| NITRFQTLLALHRSY | SPIKE | 0.66 | Probable antigen | NON-ALLERGEN | 100% |
| NNLDSKVGG | SPIKE | 0.73 | Probable antigen | NON-ALLERGEN | 100% |
| QIPFAMQMAYRFNGI | SPIKE | 0.64 | Probable antigen | NON-ALLERGEN | 100% |
| QKFNGLTVLPPLLTD | SPIKE | 0.62 | Probable antigen | NON-ALLERGEN | 100% |
| QLTPTWRVY | SPIKE | 0.74 | Probable antigen | NON-ALLERGEN | 100% |
| QSLLIVNNATNVVIK | SPIKE | 0.67 | Probable antigen | NON-ALLERGEN | 100% |
| RDIADTTDAVRDPQ | SPIKE | 0.59 | Probable antigen | NON-ALLERGEN | 100% |
| RTQLPPAYTNS | SPIKE | 0.69 | Probable antigen | NON-ALLERGEN | 100% |
| RVYST | SPIKE | 0.83 | Probable antigen | NON-ALLERGEN | 100% |
| SGTNGTKRFDN | SPIKE | 0.71 | Probable antigen | NON-ALLERGEN | 100% |
| SQSIIAYTMSLGAEN | SPIKE | 0.64 | Probable antigen | NON-ALLERGEN | 100% |
| TRFQTLLALHRSYLT | SPIKE | 0.68 | Probable antigen | NON-ALLERGEN | 100% |
| VITPGTNTSN | SPIKE | 0.7 | Probable antigen | NON-ALLERGEN | 100% |
| VLPFNDGVY | SPIKE | 0.7 | Probable antigen | NON-ALLERGEN | 100% |
| VLSFELLHAPATVCG | SPIKE | 0.58 | Probable antigen | NON-ALLERGEN | 100% |
| VRQIAPGQTGKIAD | SPIKE | 0.59 | Probable antigen | NON-ALLERGEN | 100% |
| VVLSFELLHAPATVC | SPIKE | 0.54 | Probable antigen | NON-ALLERGEN | 100% |
| VVVLSFELLHAPATV | SPIKE | 0.58 | Probable antigen | NON-ALLERGEN | 100% |
| YGFQPTNGVGYQ | SPIKE | 0.67 | Probable antigen | NON-ALLERGEN | 100% |
| YKLPDD | SPIKE | 0.79 | Probable antigen | NON-ALLERGEN | 100% |
| YQAGSTPCNGV | SPIKE | 0.66 | Probable antigen | NON-ALLERGEN | 100% |
| IGAVILRGHLRIAGH | MEMBRANE | 0.61 | Probable antigen | NON-ALLERGEN | 100% |
| ILRGHLRIAGHHLGR | MEMBRANE | 0.57 | Probable antigen | NON-ALLERGEN | 100% |
| LNTDHSSSSD | MEMBRANE | 0.75 | Probable antigen | NON-ALLERGEN | 100% |
| LSYYKLGASQRVAGD | MEMBRANE | 0.58 | Probable antigen | NON-ALLERGEN | 100% |
| SYYKLGASQRVAGDS | MEMBRANE | 0.59 | Probable antigen | NON-ALLERGEN | 100% |
| VIGAVILRGHLRIAG | MEMBRANE | 0.58 | Probable antigen | NON-ALLERGEN | 100% |
| DQIGYYRRATRRIRG | NUCLEOCAPSID | 0.61 | Probable antigen | NON-ALLERGEN | 100% |
| QIGYYRRATRRIRGG | NUCLEOCAPSID | 0.62 | Probable antigen | NON-ALLERGEN | 100% |
| RIRGGDGKMKDL | NUCLEOCAPSID | 0.61 | Probable antigen | NON-ALLERGEN | 100% |
| AAGTTQTACTDDN | ORF 1a | 0.67 | Probable antigen | NON-ALLERGEN | 100% |
| AAIFYLITPVHVMSK | ORF 1a | 0.66 | Probable antigen | NON-ALLERGEN | 100% |
| AARYMRSLKVPATVS | ORF 1a | 0.63 | Probable antigen | NON-ALLERGEN | 100% |
| AIILASFSASTSAFV | ORF 1a | 0.65 | Probable antigen | NON-ALLERGEN | 100% |
| ASFNYLKSPNFSKLI | ORF 1a | 0.64 | Probable antigen | NON-ALLERGEN | 100% |
| AVVLLILMTARTVYD | ORF 1a | 0.62 | Probable antigen | NON-ALLERGEN | 100% |
| EAARYMRSLKVPATV | ORF 1a | 0.58 | Probable antigen | NON-ALLERGEN | 100% |
| FLCLFLLPSLATVAY | ORF 1a | 0.63 | Probable antigen | NON-ALLERGEN | 100% |
| FMRFRRAFGEYSHVV | ORF 1a | 0.58 | Probable antigen | NON-ALLERGEN | 100% |
| IINLVQMAPISAMVR | ORF 1a | 0.6 | Probable antigen | NON-ALLERGEN | 100% |
| KEMYLKLRSDVLLPL | ORF 1a | 0.58 | Probable antigen | NON-ALLERGEN | 100% |
| KSAFYILPSIISNEK | ORF 1a | 0.61 | Probable antigen | NON-ALLERGEN | 100% |
| KYKFVRIQPGQTFSV | ORF 1a | 0.61 | Probable antigen | NON-ALLERGEN | 100% |
| LGSLIYSTAALGVLM | ORF 1a | 0.55 | Probable antigen | NON-ALLERGEN | 100% |
| LVNKFLALCADSIII | ORF 1a | 0.65 | Probable antigen | NON-ALLERGEN | 100% |
| NHNFLVQAGNVQLRV | ORF 1a | 0.65 | Probable antigen | NON-ALLERGEN | 100% |
| PLYAFASEAARVVRS | ORF 1a | 0.57 | Probable antigen | NON-ALLERGEN | 100% |
| RGMVLGSLAATVRLQ | ORF 1a | 0.61 | Probable antigen | NON-ALLERGEN | 100% |
| RVLGLKTLATHGLAA | ORF 1a | 0.56 | Probable antigen | NON-ALLERGEN | 100% |
| SRVLGLKTLATHGLA | ORF 1a | 0.58 | Probable antigen | NON-ALLERGEN | 100% |
| STQEFRYMNSQGLLP | ORF 1a | 0.59 | Probable antigen | NON-ALLERGEN | 100% |
| TRYVLMDGSIIQFPN | ORF 1a | 0.6 | Probable antigen | NON-ALLERGEN | 100% |
| YLYFIKGLNNLNRGM | ORF 1a | 0.62 | Probable antigen | NON-ALLERGEN | 100% |
| KKRWQLALSKGVHFV | ORF 3a | 0.63 | Probable antigen | NON-ALLERGEN | 100% |
| KRWQLALSKGVHFVC | ORF 3a | 0.59 | Probable antigen | NON-ALLERGEN | 100% |
| VKHVYQLRARSVSPK | ORF 7a | 0.57 | Probable antigen | NON-ALLERGEN | 100% |
